# Understanding pollination in urban food production: the importance of data validation and participant feedback for citizen science project design

**DOI:** 10.1101/2024.11.09.622771

**Authors:** Elizabeth Nicholls, Leah Salm, Maria Clara Castellanos, Parthib Basu, Soumik Chaterjee, Adrian Ely, Helena Howe, Dave Goulson

## Abstract

There is a significant knowledge gap regarding the pollination needs of urban farming, partly due to barriers for researchers in accessing these growing spaces. Involving growers in data collection offers a potential solution but presents other challenges in terms of data accuracy and participant retention.
We developed a citizen science methodology for monitoring plant-pollinator interactions in urban food systems and evaluated the accuracy of data collected by growers by comparison to data collected by a professionally trained researcher. We also collected feedback from participants at the mid- and endpoints of the project regarding their experiences of taking part.
While there was some agreement between the datasets in terms of the crops most (raspberries and squash) and least attractive (tomatoes) to insects, relying only on the dataset collected by growers themselves (citizen scientists) would lead to an overestimation of the generality of relationships between crops and pollinating insects in urban food production. Possible reasons for discrepancies between the datasets include species misidentification and non-reporting of surveys where no insects were observed by citizen scientists.
Citizen scientists reported lack of time, concerns about data accuracy and too complex methods as barriers to participation. Implementation of their suggestions for improvements led to a 66% increase in participation in the following year, demonstrating the importance of maintaining a two-way dialogue between participants and project organisers. Citizen scientists also reported an increased appreciation and understanding of insect pollinations following participation, highlighting an additional benefit of involving urban growers in data collection.

**Social Impact Statement:** Growing food in urban areas can contribute towards sustainable food production. We know little about urban crop pollination, and data collection can be challenging due to restricted access to growing spaces. We developed and tested the effectiveness of a method for urban growers themselves to collect data on insects visiting their crops. Comparing data to that collected by a professional researcher, we found similarities in terms of the most and least attractive crops to pollinators, but there were also large discrepancies in the datasets that mean there are limitations to relying on grower-collected data alone to understand urban plant-pollinator interactions.

## Introduction

Urban farms are estimated to account for close to 6% of all cropped areas worldwide (Thebo et al. 2014) and while often small-scale and labour-intensive, have been shown to be as productive as conventional rural farms (Altieri 2009; Laughton 2017; McDougall et al. 2019; Edmondson et al. 2020; Nicholls et al. 2020; Dobson et al. 2021a). Typically less reliant on synthetic inputs such as fertilisers and pesticides, urban farming is increasingly recognised as a potential solution to sustainably meeting some of the food demands of an urbanising global population (Nicholls et al. 2020). While substantially fewer studies have investigated the impacts of urban food growing on wildlife compared to studies of conventional agricultural systems, available research demonstrates benefits to several taxonomic groups (Colding et al. 2006; Matthies et al. 2015; Borysiak et al. 2017) including birds and insects (Quesada and MacGregor-Fors, 2010; Lin et al. 2015). For example, a large-scale study of pollinator diversity and abundance across four UK cities found that pollinating insects such as bees are more diverse and abundant in urban growing areas compared to other land uses, including nature reserves (Baldock et al. 2019). While approximately one third of food grown under conventional farming systems is estimated to be dependent on insect pollination (Aizen et al. 2008), a significant knowledge gap remains regarding the pollination needs of crops commonly grown in urban areas, and whether pollinator populations are sufficiently large and diverse to sustain or even expand urban food production (but see Bennett and Lovell 2019; Nicholls et al. 2023; Cohen et al. 2021).

There are several barriers to accurately quantifying the productivity and pollination needs of urban growing spaces, including the frequently *ad hoc* nature of urban farms, and uncertainties concerning land ownership and access (Holligan and Howe 2024), particularly in low and middle-income countries where urban farming is more widespread (Zezza and Tasciotti 2008; Hamilton et al. 2013; Thebo et al. 2014). Citizen science, defined as ‘scientific work undertaken by members of the general public, often in collaboration with or under the direction of professional scientists and scientific institutions’ (Oxford English Dictionary, 2012) may offer a potential solution. As well as permitting data collection over larger spatial areas and longer time scales than can typically be achieved by professional research teams, at lower cost (Encarnação et al. 2021; Parretti et al. 2023) an additional benefit of involving non-professionals in data collection is the ability to access remote or restricted spaces, such as private gardens and allotments in the urban farming context. Citizen science may also be referred to as community science, participatory action research (PAR) or public participation in scientific research (PPSR) depending on the academic field.

There is growing interest in using citizen science to contribute to monitoring the 17 UN-agreed Sustainable Development Goal (SDG) targets and indicators (Fritz et al. 2019; Cappa et al. 2022), with a recent review suggesting that citizen scientist-generated data has the greatest potential to contribute to monitoring the goals ‘11: Sustainable cities and communities’ and ‘15: Life on land’ (Fraisl et al. 2020). While examining the contribution of small-scale food production in urban areas to the SDGs, Nicholls et al. (2020)identified several synergies between these two goals. Citizen science has already been used to estimate crop yields and characterise farming practices in urban contexts by a number of researchers, including ourselves (McDougall et al. 2019; Nicholls et al. 2020; Dobson et al. 2021b). There are also numerous citizen science schemes for monitoring pollinators and, to a lesser extent, plant-pollinator interactions (Kremen et al. 2011; Domroese and Johnson 2017; Kleinke et al. 2018; Falk et al. 2019; Garratt et al. 2019a; Mason and Arathi 2019; Bloom and Crowder 2020; Serret et al. 2022). A recent meta-analysis by Koffler et al. (2021) suggests that progress on five out of the 17 SDGs could be addressed by studying bees.

As well as generating data, citizen science can also lead to behavioural change (Dunkley 2017; Dickinson et al. 2012; Wals et al. 2014; Peter et al. 2021;but see Druschke and Seltze 2012). For example, a survey of 139 participants in butterfly-related citizen science projects in the US found that after participating in such projects, 40% of respondents had reduced herbicide and insecticide use and 50% had planted nectar-rich or host plants for butterflies (Lewandowski and Oberhauser 2017). Behaviour change arising from participation in citizen science projects could be particularly beneficial in urban areas where there is high potential for the adoption of pro-environmental actions in private gardens or growing spaces to benefit biodiversity (Cosquer et al. 2012; Lewandowski and Oberhauser 2017; Griffiths-Lee et al. 2020; Sturm et al. 2021; Toomey and Domroese 2013)

A major barrier to the adoption of citizen or community science data in monitoring schemes are the widely debated concerns regarding the quality and accuracy of data collected by non-professionals (Kosmala et al. 2016; Lukyanenko et al. 2020). Protocols need to be simple enough for non-specialists to perform, which can limit the types of data and/or level of detail that can be extracted, and even with a simple protocol, there may still be concerns regarding the accuracy or reliability of data collected by non-scientists. There are a range of options available to maximise the quality of data, however, from direct comparison with professionally collected data or expert validation of data submitted by citizen scientists to automated filtering of outliers (Kosmala et al. 2016). By adopting one or more of these quality controls, several studies have shown that, given appropriate training and feedback, biodiversity data collected by community scientists can be on par with data collected by professional scientists (Kremen et al. 2011; Kosmala et al. 2016).

Here our aim, as an inter-disciplinary team of researchers, was to develop a citizen science methodology for monitoring plant-pollinator interactions in urban food systems and to evaluate the accuracy of data collected by growers by comparing it to data collected by a professionally trained researcher using an identical methodology. We focus on a comparison of the similarity of plant-pollinator interaction data collected by a professional researcher versus citizen scientists: a more in-depth analysis and discussion of the implications of the findings in the context of urban crop pollination can be found in Nicholls et al. (2023).

Given that the project was initiated by a team of professional researchers (top-down organisation), and a major project goal was to develop a scientific method to answer research questions rather than to solve a particular community issue, we have defined this as a citizen science rather than a community science project (Lin-Hunter et al. 2023). However, though initiated by researchers, we collected feedback from participants at the mid and endpoints of the project, and where possible incorporated this feedback into the methodology for the second year of the study. The aim was to improve participation in the project, given that a major challenge of citizen science is the often-high levels of non-engagement from recruited and trained participants that may arise for several reasons, including methods that are too difficult or time-consuming (Birkin and Goulson 2015; Kleinke et al. 2018; Bloom and Crowder 2020). We also asked participants for feedback on the research aims and questions and suggestions regarding priorities for future research directions in the broad area of urban food production. At the end of the project, we asked participants to self-evaluate how their ability to identify insects (knowledge gain) and gardening practices (behaviour change) were affected by participation in the project, if at all. By maintaining a two-way dialogue between scientists and participants throughout the project, our work provides important insights into the process of designing and delivering a citizen science project that we expect will be of broad interest and value to academics beyond the fields of plant and pollinator biology, since many of the considerations we raise here will be of relevance to various types of citizen science project.

## Materials and Methods

### Study location and volunteer recruitment

The study was conducted in the city of Brighton & Hove, Southeast England, UK between 2017 and 2019. Brighton has a population of 273,369 (UK census 2011) and according to the city allotment strategy published in 2014, there are 6,000 allotment growers (∼2% population), farming 3,100 plots over 37 allotment sites. In total, allotments (a plot of land rented by an individual or community group for horticulture) cover an area of approximately 50 ha, and an unknown proportion of home gardens are also used to produce food in the city.

During the two years (April 2017- September 2018) the project was active, we recruited 184 allotment-holders and home-growers across the city to participate in ‘Team PollinATE’. In Year 1 (2017) we restricted participation to those who grew food on an individual or community allotment plot (n=77 citizen scientists), however in Year 2 (2018) participation was also opened to those who grew food in a private garden (n=130 recruits total). Twenty-three people took part in both years of the study. We promoted the project *via* posters placed each year at allotment sites across the city, emailing lists (Brighton & Hove Allotment Federation (BHAF); Brighton & Hove Food Partnership, The Living Coast), social media and community gardening-themed events such as Seedy Sunday and Brighton & Hove Organic Gardening Group (BHOGG) workday. Collaboration with BHAF was particularly critical in raising awareness of the project, and in gaining access to sites for surveying by the professional scientist. Participants signed up *via* the project website (https://www.thebuzzclub.uk/team-pollinate), by emailing the lead organiser or by using a paper-based sign-up form at community events. Once signed up, participants received a welcome pack (Additional File S1) in the post and were added to an e-mailing list to receive monthly updates.

### Citizen scientist data collection

We asked participants to collect data on the types of pollinating insects visiting their flowering food crops throughout the growing season. Participants were provided with recording sheets and instructions on how to perform a survey, as well as pollinator-identification materials (Additional File S1). They were invited to attend an optional in-person training session during which they were given a talk about the aims of Team PollinATE, and how to identify different types of pollinator.

We asked participants to perform up to two pollinator surveys per month between April and September, or for as long as they had insect-pollinated crops flowering in their growing space. They were asked to perform an instantaneous count of the number of pollinators observed when looking at individual flowers of a particular crop type one by one in sequence until all flowers had been observed (Levin et al. 1968;Vaissière et al. 2011). For more details on the pollinator surveying method guidance see Methods S1. Participants were also asked to collect data on any pest issues associated with flowering crops in their plots, as well as the weight of yields harvested from their flowering crops each year. These methods and data are published in Nicholls et al. (2020).

### Researcher data collection

To validate the accuracy of the citizen scientist-collected data, we obtained permission from a subset of participants for a professionally trained scientist to visit their growing space and conduct pollinator surveys using the same method over the same period. Plots were distributed across nine allotment sites (1-29 plots per site, Table S1) located in different regions of the city (Fig. 1, orange areas). Of those participants who permitted their plots to be surveyed, 13 submitted their data on pollinator visits as part of the citizen science component of the study.

**Figure 1.**
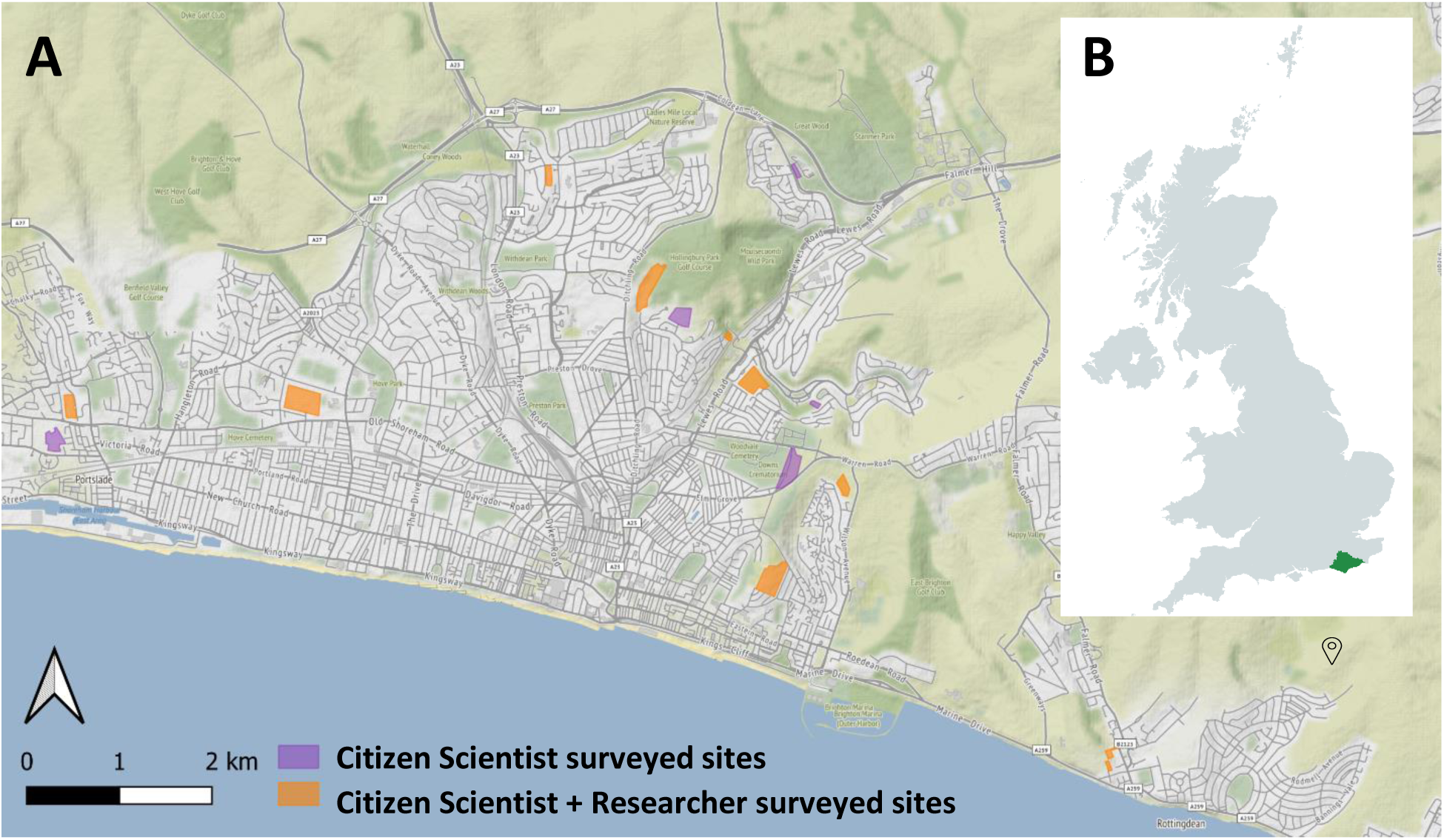
a) Location of allotment sites surveyed by citizen scientists and one researcher for pollinators across the city of Brighton & Hove (orange, n=9) and allotment sites surveyed by citizen scientists only (purple, n=6) and b) study location the city of Brighton & Hove (black marker) in East Sussex (green highlighted area), UK. For a full list of sites and the number of citizen scientists submitting data from each allotment site see Table S1. UK chart created in mapchart.net.

### Participant feedback and results dissemination

At the mid-point and endpoint of the project we held a ‘Feedback Event’ for the participants where we presented a summary of the data that had been submitted and some of the preliminary conclusions. At the mid-point event, participants were asked to give feedback on their experience of taking part and how the methodology and experience could be improved. Feedback at the mid-point event was collected using several approaches; 1) sticker voting on a set of five questions (Table S2), comment boards under four questions regarding their experience of taking part in Team PollinATE and how we could improve the methodology which attendees could add to (Table S3) and 3) a ‘World Café’ where attendees circulated between two tables hosted by a member of the research team who led guided discussions on the broad topics of i) participants’ experience of taking part in Team PollinATE ii) suggestions for methodological improvements and ii) participants’ perceptions of the research aims and questions being investigated by Team PollinATE (Table S4). Where possible, the participants’ suggestions were incorporated into the methodology for the second year of the study. We also asked participants what they thought about the scientific aims of the project.

At the endpoint event we used a paper-based survey (also available online) (Additional File S1) to collect data and asked participants to estimate their ability to identify pollinating insects before and after taking part in the study, and whether their gardening habits had changed (DataSet S1). To further disseminate the results, we produced a leaflet summarising the aims of Team PollinATE, featuring an infographic to visualise the main results of the project in a simplified and accessible manner (Additional File S1).

### Data Analysis

#### Analysis of crop-pollinator data

All quantitative analysis was conducted in R v. 4.2.1 [R Core Team, 2022]. Before analysis, we inspected the data and removed any observations recorded for crops or flowering plants that were not insect-pollinated. There were a limited number of surveys for certain crops, so to avoid having low sample numbers which may affect the estimation of crop-pollinator network structure (Fründ et al. 2016), crop types were condensed into ten categories as in Nicholls et al. (2023) (Methods S2).

To compare the distribution of researcher (R) and citizen scientist (CS) surveys (Data Set S2) across crops and sampling months, we used z-tests to conduct pairwise comparisons of the proportions of observations, with Bonferroni corrections for multiple comparisons. To compare the total number of insect visits observed between R and CS datasets across different sampling months and crops, we ran a generalised linear model (GLM), with a negative binomial distribution to handle overdispersion in the data, using package *‘lme4’*. We used the formula glmer.nb (Total Insects Observed ∼ Observer Type * Crop Type + Observer Type * Sampling Month), to test the effects of the main factors and their interactions. Model fit was checked using residual diagnostics via the package *‘DHARMa’* (Hartig 2018).

To visualise the similarities in crop visitation between pollinator groups in data collected by the researcher and citizen scientists we used non-metric multidimensional scaling (NMDS) ordination, using Bray–Curtis dissimilarities of the relative visitation frequencies. We also compared plant-pollinator network metrics, using similar methods to Nicholls et al. (2023) (Methods S2). Ordination is used to reduce the complexity of large datasets and to visualize how communities or species differ from each other across multiple dimensions. Network analysis, on the other hand, focuses on the interactions between species, revealing how they are connected i.e. who interacts with whom and how frequently and can highlight key species or relationships that play a critical role in maintaining community stability. Together, these methods provide a deeper understanding of how species interact and community dynamics, offering insights that simple numerical comparisons might miss. We constructed three separate networks from the following data sets i) all researcher-collected data (R), ii) all citizen scientist-collected data (CS) and iii) citizen scientist-collected data matched by allotment site to the researcher-collected data (CSM). Given that network metrics can be affected by factors such as the number of species in the network and sampling effort (Fründ et al. 2016), we calculated the means from 1,000 null models using the Patefield algorithm (Patefield 1981) which fixes the network size while shuffling interactions randomly. We then used z-scores ((observed value-null mean)/null standard error) to test for significant differences from the null model distribution (Dormann 2022).

#### Analysis of participant feedback

The feedback data from participants collected at the mid- and end-point events was analysed through descriptive analysis for the categorical questions in the paper-based survey, and by thematic analysis for the open-ended questions, ‘World Cafe’ discussions and comment board suggestions. Themes related to areas of barriers to participation and suggestions for improvements in the future and arose inductively based on the specific context of urban food growing and pollinator interactions. Two researchers were involved in this thematic coding process. Key quotes which best described the nuance of the themes were extracted from the transcripts of the ‘World Cafe’ discussions and responses to the open-ended survey questions

## Results and Discussion

### Frequency and timing of crop surveys

Over the two years, 33 citizen scientists (CS) submitted data (18% of volunteers recruited), and conducted 580 crop surveys across 21 different sites, including individual (n= 12 sites) and community allotment sites (n= 3) and private home gardens (n= 6) across the city of Brighton & Hove (Fig. 1, purple and orange areas), recording a total of 1,552 insect visits. The researcher (R) conducted a total of 1,579 crop surveys, in 83 plots (individual=78, community = 5) across nine allotment sites (Fig. 1, orange areas), recording 1,923 insect visits to flowering crops. The number of surveys per crop type, and the number plants of each type surveyed at a particular plot were variable within both data sets (Table S5) due to individual preferences in the types and numbers of each crop grown by each plot-holder. The proportion of total surveys conducted across the crop types also differed slightly between datasets, CS had fewer observations of apples (z=-3.220, p=0.013) and more observations of tomatoes (z=3.810, p<0.001; Table S6).

The timing of the crop surveys during the day was very similar, with just over one-third of surveys taking place in the morning (R= 35%, CS= 39% observations). However, the proportion of total surveys conducted within each month was significantly different between the R and CS datasets, with CS conducting proportionally more surveys in June (z=5.535, p<0.001) and July (z=2.790, p=0.032) and fewer in August (z=-6.637, p<0.001) (Table S7-8).

CS recorded significantly more insects per survey compared to R in all months except April (χ^2^= 17.342, df=5, p=0.004). For R the mean number of insects per crop survey peaked in July (mean ±SD= 1.54±3.46). In contrast, the number of observations per survey peaked in September for CS (Fig. 2; 3.53±8.99), at the end of the growing season, which was the month that R observed the lowest number of insects per survey (0.77±1.52). However, this finding is likely driven by an extreme outlier, given one citizen scientist recorded seven observations of each insect type except social wasps visiting squash flowers (n= 49 insects for a single survey). The consistency in the value across insect categories suggests this may have been an error in electronic data entry. If this entry is removed, then the number of observations per survey peaks in August for the CS data (3.24±5.50)

**Figure 2.**
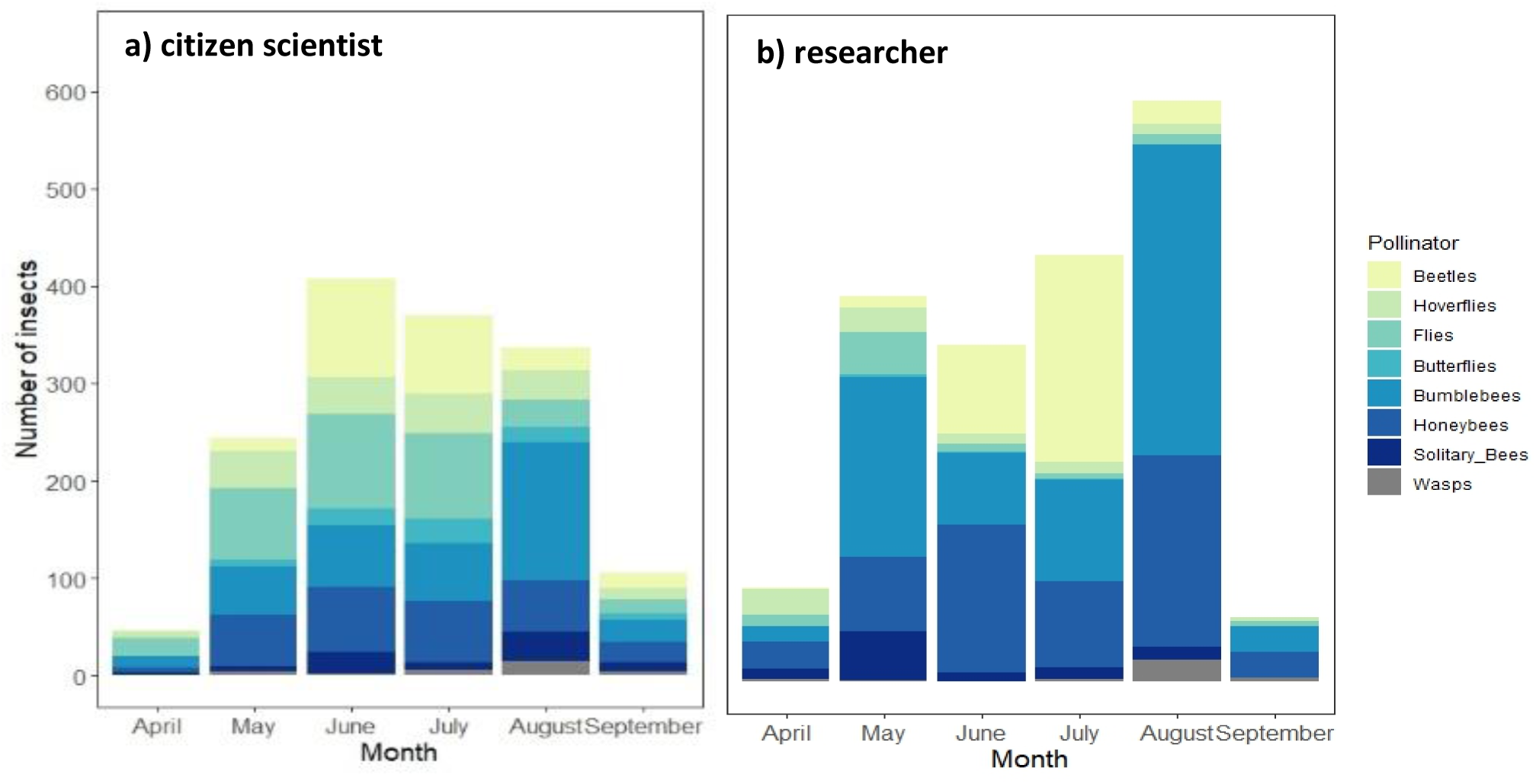
Total number of insects of each pollinator category recorded per month by a) citizen scientists and b) the researcher. Data from both years of the study combined.

### Frequency of insect observations

CS also observed significantly more insects per survey for all crops except Cherry/Plum and Cucumber (Fig. 3; χ^2^= 38.533, df=9, p<0.001; emmeans pairwise comparisons; Table S9-10). While the crop observations were intended to be instantaneous counts of insects visiting the open flowers of crops, at the feedback event participants in the current study reported feeling demotivated when they didn’t observe insects visiting their crops during a survey. Therefore, the substantially higher number of insects recorded per survey by the citizen scientists suggests that their recordings may have been biased, perhaps by waiting longer for insects to land and/or recording insect visits to parts of the plant other than the flowers. It is also possible that despite being instructed to, citizen scientists simply did not submit data from crop observations where no insects were recorded visiting the flowers. Indeed, two participants reported in the feedback survey circulated at the end of year two that they ‘*didn’t submit nil returns*’.

**Figure 3.**
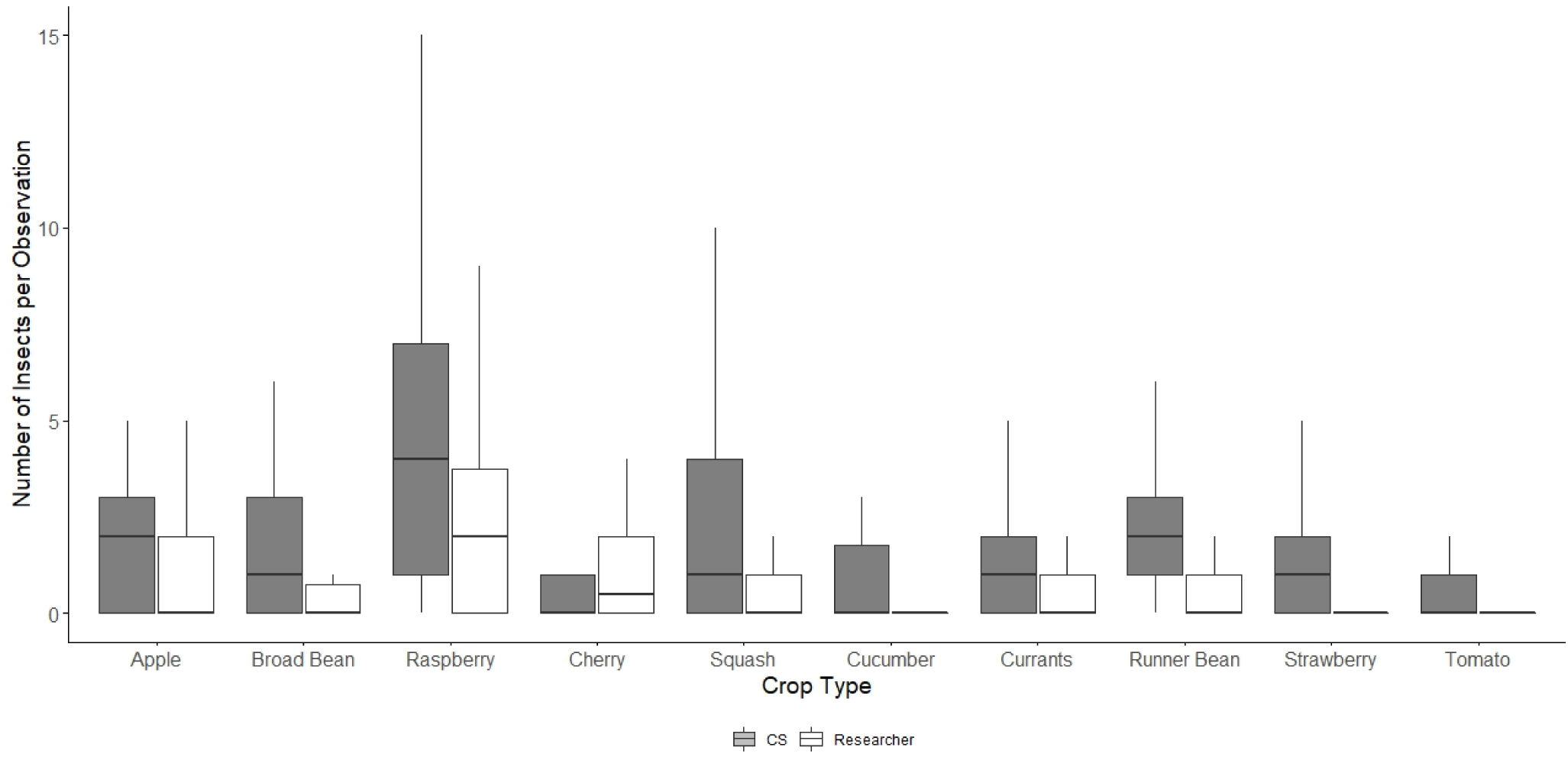
Number of insects per observation by crop for CS (grey) and researcher (white) data sets. The box limits represent the interquartile range, the line inside the box represents the median and the whiskers show the variability in upper and lower quartiles. Outliers beyond 1.5 times the interquartile range (IQR) are not displayed in this plot, to make visualisation of differences between researcher and citizen scientist data clearer.

There was some agreement between the datasets in terms of the most and least attractive crops to insect visitors. For both, raspberries were the crop with the highest number of visits per observation (Fig. 3; Table S4; R= 2.60±3.31 visits/observation; CS= 5.30±5.50), and pumpkins and squash were also very attractive (R= 1.15±2.60; CS= 2.90±5.26), with the latter driven predominantly by visits from beetles (Fig 4). R recorded the lowest number of visits per observation on tomatoes (Fig. 4; 0.06±0.23 visits/observation) and broad beans (0.37±0.72). CS also recorded low visitation rates to tomatoes (0.83±1.36) but recorded three times the number of visits to broad bean flowers, on average (1.57±1.91). This difference could potentially be explained by citizen scientists recording non-legitimate (robbing) visits by short-tongued bees since this concept was not discussed with participants.

**Figure 4.**
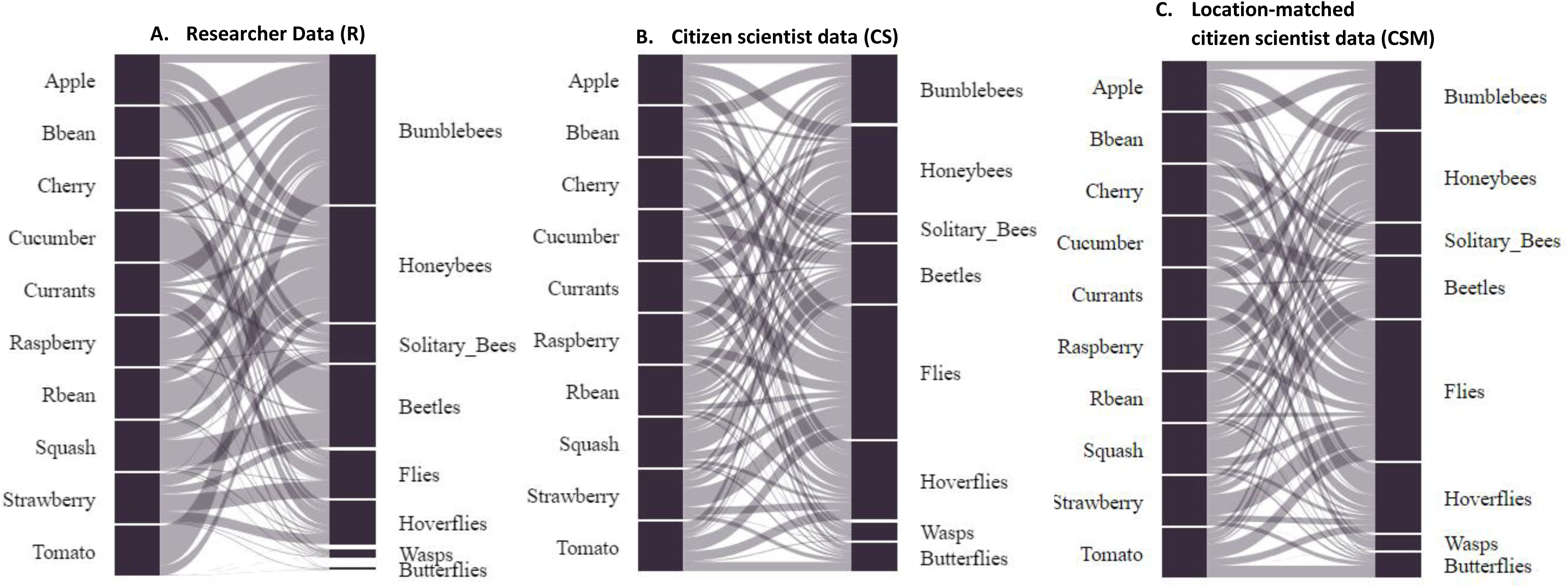
Visitation networks for insect-crop interactions based on a) all researcher-collected data (R) b) all citizen scientist-collected data (CS) and c) citizen scientist-collected data only from plots located at the sites surveyed by the researcher (CSM). Crop visits are standardised according to sampling effort by dividing the number of each insect type or species by the total number of insect visits to a particular crop to calculate the proportion of total insect visits. The size of the bar indicates the proportion of total insect visits that are represented by each insect group. Only legitimate flower visits and data for crops where there are >5 observations are included in the analysis.

There was also a major discrepancy in the number of visits observed to tree fruits such as cherries and plums, with citizen scientists recording far fewer visits to this crop type (Fig. 3; R= 1.42±2.03 visits/observation; CS= 0.82±1.63), potentially due to difficulties in observing the flowers of very large trees. When comparing researcher, farmer and volunteer collected data in apple orchards, Garratt et al. (2019) similarly found that researchers recorded significantly more blossom visits by hoverflies and solitary bees compared to volunteers.

There was also some agreement in terms of the most frequently observed insect pollinator. For both datasets, honeybees were the most frequently observed insects visiting flowering crops (Fig. 4), however the difference in the proportion of total observations of each insect group was significant for all insect groups except beetles, solitary bees and social wasps (Fig. 4, Table S11). While bumblebees were the second most frequently observed insect visitor by the researcher, accounting for just under a third of all insect visits, citizen scientists recorded a similarly high proportion of visits by flies, and bumblebees, with each insect group accounting for close to one fifth of all visits (Fig. 3).

Similarly, hoverflies accounted for a much higher proportion of flower visits in CS data compared to R (Fig. 3). These discrepancies may in part be explained by errors in the identification of the different insect groups, but it is surprising that conspicuous and charismatic groups such as honeybees and bumblebees were under-recorded. In a previous study (Griffiths-Lee et al. 2023) we also examined identification bias in pollinator-focused citizen science studies, comparing methods of insect sampling; pan traps and sticky traps with expert verification, to bee walks where insects were identified on the wing by participants only. In this study, the numbers of conspicuous insects (honeybees and bumblebees) were similar between citizen scientists and researchers, but solitary bees were under-estimated and social wasps and hoverflies were overestimated by citizen scientists. Other studies have also observed that less conspicuous insect groups are likely to be under-recorded by citizen scientists (Kremen et al. 2011; Maher et al. 2019).

### Insect-crop interactions

To visualise the similarities in crop visitation patterns between pollinators in the datasets collected by researchers and citizen scientists, we conducted a NMDS analysis. The NMDS ordination had a stress value of 0.12, indicating a fair representation of the dissimilarity in crop visitation patterns between pollinator groups. The NMDS ordination plot (Fig. 5) shows the distribution of crop visitation patterns. Observations of crop-insect interactions recorded by the researcher and citizen scientists are represented by different polygons. Though there is some overlap in polygons, the Analysis of Similarity (ANOSIM) showed a significant difference in crop visitation patterns in data collected by the researcher and citizen scientists (R= 0.155, p=0.021). An indicator species analysis revealed that butterflies (indicator value = 0.561, p=0.0017) and flies (indicator value= 0.560, p=0.013) were more frequently associated with data collected by citizen scientists. In the researcher-collected data, butterflies were recorded most infrequently, with only 5 visits to crop flowers recorded in total across 1579 crop observations (<1% total visits). However, the citizen scientists recorded 73 visits by butterflies (Fig. 4, 12.6% visits), to all crop types except tree fruit, with the highest number of visits to raspberries, squash, runner beans and strawberries. Perhaps if participants waited longer for insects to land and/or recorded insect visits to parts of the plant other than the flowers, this may explain the higher number of fly and butterfly recordings in the CS dataset, given the propensity of insects from these groups to land and rest on plant leaves, for example. Repeating the analysis comparing the researcher data (R) with citizen science data only from sites that were also surveyed by the researcher to account for localised effects on insect abundance site (CSM) did not lead to any qualitative differences compared to the initial R vs. CS comparison. The results of the ANOSIM (R= 0.147, p=0.029) and indicator species analysis were qualitatively the same as the original R vs CS comparison (Table S12).

**Figure 5.**
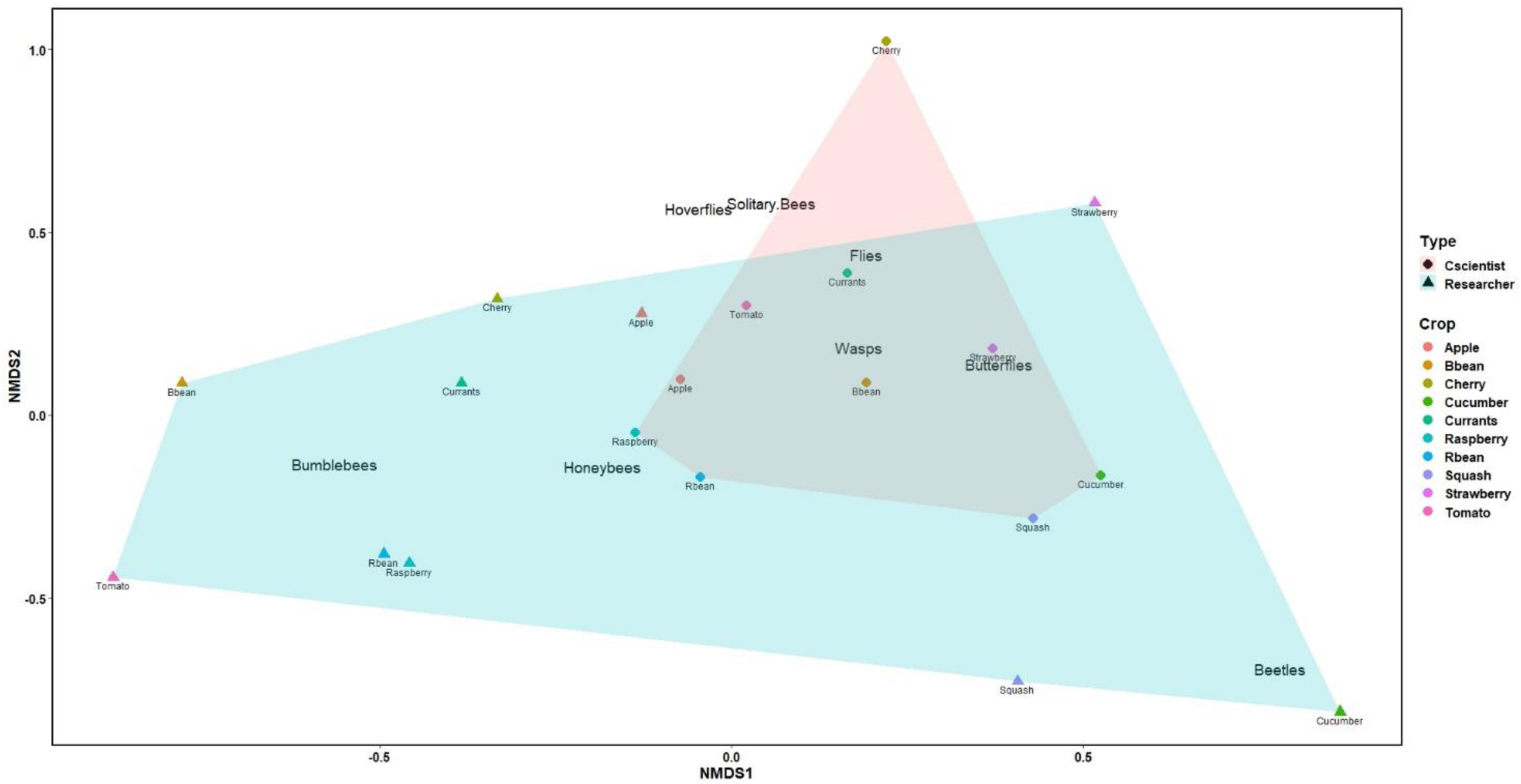
Results of nonmetric multidimensional scaling (NMDS) run with a matrix of dissimilarities (Bray–Curtis) of the relative frequency of crop visitation by the different pollinator groups (stress = 0.12) in data collected by a trained researcher (diamonds) or citizen scientists (triangles). The shorter the distance between insect pollinator taxonomic groups, the greater the overlap in crop types visited. Polygons represent the grouping of samples by dataset (researcher = blue; citizen scientist = red), with the greater the degree of overlap indicating greater similarity in visitation patterns.

To examine the frequency and strength of insect-crop interactions we constructed plant-visitor networks for both R and CS data for comparison (Fig 4a-b) and calculated network metrics of most ecological relevance to the current data set and sampling approach (Table 1). The crop-insect visitation network constructed from the researcher-collected data (Fig. 4a) was more specialised than the null model predictions, with a significantly higher *H*_2_′ value (Table 1), though the network was still fairly generalised (H’_2_=0.34) indicating that most pollinator groups interact with multiple crop types. The CS network (Fig. 4b) had a H’_2_ value of 0.16 (Table 1) and though significantly more specialised than the null model predictions (0.01), was much less specialised than the R network, indicating that citizen scientists recorded pollinator groups interacting with more crop types than the researcher. The R network had a moderate level of nestedness (WNODF= 41.08) and was significantly less nested than the null model (Table 1). This suggests that the interactions recorded by researchers are moderately nested but exhibit a non-random structure. In contrast, the CS network was less nested than the R network (WNODF= 33.54) and was not significantly different from the null model (43.81), suggesting that the observed level of nestedness in the citizen scientist data could be due to random chance.

**Table 1.** Group-level metrics for each visitation network. Sampling effort per crop type was standardised by dividing the number of visits by each insect group/species by the total number of insect visits to a particular crop, thus calculating the proportion of visits by each insect group/species. WNODF stands for Weighted Nestedness based on Overlap and Decreasing Fill. Observed values in bold are significantly different from the model predictions (p<0.001).

| Metric |  | Network |  |  |
| --- | --- | --- | --- | --- |
|  |  | R | CS | CSM |
| **Network specialization**<br>($H_2'$) | *Observed value* | **0.34** | **0.16** | **0.20** |
| | *Null mean ± SD* | 0.01±0.00 | 0.01±0.00 | 0.02±0.00 |
|  | *Corrected value* | 0.33 | 0.14 | 0.18 |
|  | *Z-score* | 153.97 | 58.21 | 62.99 |
|  | *P* | <0.001 | <0.001 | <0.001 |
| **Nestedness**<br>(WNODF) | *Observed value* | **41.08** | 33.54 | 40.53 |
| | *Null mean* | 66.93±6.18 | 43.81±9.78 | 50.40±7.94 |
|  | *Corrected value* | -25.85 | -10.28 | -9.87 |
|  | *Z-score* | -4.18 | -1.05 | -1.24 |
|  | *P* | <0.001 | 0.146 | 0.107 |
| **Interaction**<br>**Evenness** | *Observed value* | **0.80** | **0.90** | **0.89** |
| | *Null mean* | 0.73±0.00 | 0.85±0.00 | 0.84±0.00 |
|  | *Corrected value* | 0.07 | 0.06 | 0.05 |
|  | *Z-score* | 117.15 | 65.93 | 52.51 |
|  | *P* | <0.001 | <0.001 | <0.001 |
| **Visitor generality** | *Observed value* | **6.01** | **7.86** | **7.42** |
| | *Null mean* | 5.16±0.01 | 6.27±0.02 | 6.06±0.02 |
|  | *Corrected value* | 0.85 | 1.59 | 1.36 |
|  | *Z-score* | 62.01 | 70.96 | 56.50 |
|  | *P* | <0.001 | <0.001 | <0.001 |
| **Plant generality** | *Observed value* | **3.56** | **5.35** | **4.93** |
| | *Null mean* | 4.63 | 6.42±0.02 | 6.28±0.03 |
| | *Corrected value* | -1.07±0.00 | -1.07 | -1.35 |
|  | *Z-score* | -90.79 | -47.72 | -54.07 |
|  | *P* | <0.001600 | <0.001 | <0.001 |

Interactions in the R network were quite evenly distributed (Evenness= 0.80), significantly more so than the null model predictions (Table 1), suggesting that interactions were not dominated by just a few pollinator groups. Visitor generality in the R network (*i.e.* the number of interaction partners) was 6.01, and this was significantly higher than the null model predictions, indicating that insects interacted with more crops than would be expected at random. In contrast, plant generality was 3.56 which was significantly lower than the null model prediction of 4.63. Crop-insect interactions in the CS network were more evenly distributed compared to both the null model predictions and the researcher network (Evenness= 0.90), and visitor generality was higher (7.86), indicating that citizen scientists recorded insects visiting a broader range of crop types compared to the researcher. Plant generality was higher than in the researcher model (5.35), however still significantly lower than the null model prediction of 6.42. The fact that the network generated from data submitted by citizen scientists was more generalised and more even than the researcher data, with each insect group observed visiting a broader range of crops, is likely because citizen scientists made more identification errors, blurring the true pattern of insect-plant relationships.

Constructing a second network from the citizen scientist collected data (Fig. 4c, CSM), this time only including data from sites that were also surveyed by the researcher to account for localised effects on insect abundance (Table 1), slightly increased the similarity between the networks generated from researcher and citizen scientist collected data, specifically the generality (the CSM network was less generalised than the CS model) and nestedness metrics. This suggests that there is some degree of either allotment site or even plot-level effect on insect-plant interactions, which can be driven by a number of factors, such as differences in overall floral abundance and/or diversity, surrounding land-use or the proximity to semi-natural habitats (Theodorou et al. 2017; Cohen et al. 2021) which in turn may affect the delivery of pollination services to crops (Cohen et al. 2021). Given this local variation, the most appropriate comparison therefore would be between the R and CSM networks.

### Crop-visitor specialisation and species strength

There was no significant difference between R and CS networks in crop visitor specialisation (d’) (Table S13; t=1.333, df=7, p=0.164) or visitor species strength (t=0, df=7, p=1.000). In both R and CS networks, beetles had the highest crop visitation specialisation (Table S10). In the R network, this was followed by bumblebees and flies, but conversely, in the CS and CSM networks, flies had the lowest crop specialisation of any insect group. Aside from butterflies, which were recorded very infrequently in the researcher data (n=5), the next lowest crop specialisation was exhibited by the social wasps. In contrast, in the CS network, social wasps were the second most specialised crop visitor, after beetles.

In the R network, bumblebees exhibited the highest species strength, followed by honeybees, with bumblebees being the only insect group recorded visiting all crop types. While in the CS and CSM networks bumblebees and honeybees also had high species strengths, visiting up to nine of the ten different crop types, flies had the highest strength, and alongside hoverflies, were recorded visiting all crop types. Overall, each insect group had a higher number of links in the CS and CSM networks, compared to the R network.

There was a significant difference in species level specialisation (d’) between R and CS networks for crops (Table S13, t= 2.831 df= 9, p= 0.020), but when the citizen scientist data was matched to the researcher data by site, the difference in crop specialisation between R and CSM was no longer significant (t=1.916, df=9, p=0.091). There was no significant difference in crop species strength between the R and CS model (t= 0.204, df=9, p=0.850). The researcher observed a broad number of insect groups visiting cherries and plums, but in the CS network, this was the most specialised crop with the fewest interaction link. In the data collected by the researcher, cucumbers, strawberries and tomatoes were the most specialised in terms of insect visitors, and the difference with the CS network was particularly extreme for tomatoes, in which eight different insect groups were observed visiting compared to just two (bumblebees and honeybees) in the R network. Even in the case of the researcher data, it is known that honeybees are less effective at pollinating tomatoes compared to bumblebees that are capable of buzz pollination (Cooley and Vallejo-Marín 2021). This highlights a major drawback of only recording insect visits - these do not indicate the degree of pollination efficacy and thus the importance of the insect for pollination is not possible to determine with accuracy. Despite this limitation, previous pollination research has often shown that the more frequent insect visitors are also typically the most efficient pollinators (Rader et al. 2016; Ballantyne et al. 2017), so visitation data can serve as a useful proxy, particularly as a first step in identifying the most common insect visitors to food crops grown in urban areas. This type of data may be most useful in low and middle-income countries where urban growing is particularly prevalent, and provides an important source of income, but where data on urban insect-crop visitation is even more sparse than for the UK. Nonetheless, it is clear from the current study that relying only on citizen scientist data collected using the methodology described here would lead to an overestimation of the generality of relationships between crops and pollinating insects in urban food production, at least in a UK context.

### Participant feedback and real-time alterations to methods in Year 2

In Year 1, participants were asked to give feedback on their experience of taking part in Team PollinATE at the mid-point event (n= 19 attendees, ten of whom had submitted data). Voting on five set questions revealed that 100% of respondents (N=8) felt the level of communication from the project organisers in the first year of the project was just right. Just over half of respondents rated the Year 1 methods ‘Very Easy’ with one third rating them ‘Moderately Easy and 11% rating them ‘Moderately Hard’ (Table S2). However, respondents were split in terms of how difficult it was to submit data, with just under 30% of respondents finding it ‘Moderately Hard’, 30% finding it ‘Moderately Easy’ and 40% finding it ‘Very Easy’. The feedback from the comment boards indicated that participants in the first year most enjoyed learning about pollinators and becoming more aware and observant (Table S3).

In terms of barriers to participation and the submission of data, participants found that it was difficult to count the flowers in the plot, leading in some cases to concerns regarding data accuracy:

> *“I just kind of started to look [at number of flowers] and then I just kind of just felt like the inaccuracy made it a bit not worth doing”* (Citizen science participant, Year 1)

Another commonly reported barrier were the other demands at the allotment leading to a lack of time to collect data:

> *“I felt it was really time consuming. And when I go, I’ve already planned out exactly what I’m going to do. There’s already a long list, to-do stuff”* (Citizen science participant, Year 1)

Some also felt that data collection and submission methods were too difficult or demanding:

> *“I just really don’t want to go on a computer when I’m somebody that works on a computer so the idea of going on and putting data in just makes me feel oh no - I think that’s just a personal thing”* (Citizen science participant, Year 1)

These reported barriers (see Table S3-4) reflect the findings of other citizen science project studies and evaluations, where a lack of time and protocols that are too demanding or complex have been found to affect participation and data submission (Birkin and Goulson 2015; Bloom and Crowder 2020; MacPhail and Colla 2020; Serret et al. 2022).

Table 2 summarises the suggestions for improvements we received from participants and how we responded to these. In Year 2 we held more practical, in-person training events at the start of the growing season, to demonstrate the methods involved in taking part. We simplified the methods for collecting data on pollinators and provided categories for estimating the approximate number of plants to counter concerns regarding the effort and accuracy of trying to count this. We reduced the frequency of surveys to a minimum of once per month to account for people struggling for time at the allotment with all the other tasks they need to complete. We also made participation in the different parts of the project optional; participants could just count pollinators, measure harvests or record pests, or take part in all three. We made uploading information via the online form simpler, so that participants no longer had to repeat the submission of data (such as size of their growing space) each time they submitted data, instead we used their email addresses to match data across submissions. We also provided a stamped, addressed envelope for those who preferred not to use a computer to input data and offered the option of emailing us a photo or scan of their data collection sheets. We updated the project website and made an online community forum where participants could share photos or ask questions. After implementing these changes, we observed a 66% increase in data submissions between Year 1 and Year 2, highlighting the importance of piloting new citizen science methodologies and maintaining a two-way dialogue between project organisers and participants. Interestingly not all of the suggested improvements were utilised after being implemented, for example the community forum received only two posts over the course of the Year 2 growing season. A lack of time remained a frequently reported barrier to participation in Year 2, despite streamlining of the methods.

**Table 2.** The key barriers to participation identified by citizen scientists at the mid-way point of the project and how these were addressed by changes to the methodology and project organisation for Year 2.

| Key barriers to participation in Year 1 | Participant feedback examples | Changes to methodology for Year 2 |
| --- | --- | --- |
| Data accuracy concerns | *'Bothered me how inaccurate the data was- e.g. impossible to count raspberry flowers'* | Datasheets edited to include explicit categories for estimating the approximate number of plants for each crop type. |
| Support for data collection and submission | *'Going on computer off-putting'* | Provided stamped addressed envelopes for participants to post data sheets back at the end of the season. Option of emailing scans and/or photos of datasheets instead of uploading data online. |
|  | *'Reporting survey results and crop yields was fiddly- would be good to have individual account to save entering [personal] details each time'* | Participant data matched according to email address and input of site details no longer required for each submission. |
|  | *'Is there somewhere to upload photos so we can check we're doing it right e.g. community forum?'* | Added an online forum to project website where participants could upload photos and ask questions. |
| Lack of data | *'Not counting insects was very discouraging'* | Monthly email updates containing summaries of data collected in the previous month to help convey to participants that data submitted by individuals is part of a bigger project at the city scale. |
| Lack of time | *'Time! Or rather prioritising data collection over weeding, watering, planting etc.'* | Reduced frequency of surveys to a minimum of once per month. Made different parts of the project optional; participants could choose whether to count pollinators, weigh harvests or record pest control methods or a combination. |
| Participant motivation and retention | *'Possibly a more flexible survey schedule- depressing when got behind!'* |  |
| Methods too hard | *'Hard- couldn't estimate flower numbers'* | In person, practical training sessions on allotment sites. |

As well as feedback on participants’ experiences of taking part in the project in Year 1, we also asked for their feedback on the three main aims of the project, which were to 1) understand which insects pollinate which crops in urban areas 2) quantify the production of food in urban areas and 3) determine the most common pests in urban areas and the methods used to control them.

Participants felt that the data was a good baseline, and that long term data is important for observing changes over time. Many participants raised the issue of an initial learning curve and feeling overwhelmed at first, but also acknowledged that the data collection process got easier with time. Some would also seek help or reassurance from fellow growers, an advantage of a very localised project:

> *“At first you think like huh?, and then you read it again and go okay, I get that and then you do it, and then, you know…” (Citizen scientist participant, Year 1)*

> *“So it’s almost like it should come with a warning sticker. It might seem a bit of a head spin for a couple of weeks. Give yourself the chance to, you know, learn what you’re doing or something and to get used to it.” (Citizen scientist participant, Year 1)*

In response to this prompt, participants again raised concerns regarding the accuracy and the implications for the meaningfulness of the data:

> *“Once we’ve missed a two weeks slot we were like oh it’s all gone horribly wrong. Is it useful just to have snapshots spread out over the summer and especially when it’s very variable?” (Citizen scientist participant, Year 1)*

Collectively, this feedback highlighted the need to provide reassurance that even small amounts of data were useful to the project, and to provide more regular updates of data that had been collected across participants. A study of Dutch volunteer biodiversity recorders also found that volunteers have high expectations regarding the impact of the data they collect (Ganzevoort et al. 2017). For individual participants who are not familiar with ecological data collection and the importance of the absence of observations (i.e. zeroes), the perceived lack of data could be very demotivating when considered in isolation:

> *“Actually I did find, when I did get around to looking there were hardly ever any bloody bees - I would be there wandering my allotment and like oh there is a bee over there and by the time I get there the bee would have pushed off” (Citizen scientist participant, Year 1)*

Participants had many suggestions for future methodological improvements that were beyond the scope of the current study, such as asking growers to estimate the area of their plot cultivated with food versus other plants, and to test the effectiveness of pesticides in terms of the volumes applied by growers (see Table S3-4). They also had suggestions for additional areas of related research, such as the impact of environmental effects e.g. soil type, proximity to the sea on pest and pollinator abundance in urban growing spaces, the role of nocturnal pollinators, and the influence of growing bee friendly flowers on crop visitation by pollinators (see Griffiths-Lee et al. 2020).

> *“Why are we just recording for consumable crops? Because obviously, there’s so much more going on with the non-consumable commercial flowers basically. So, are they attracting also that same portion of bees into a pollinating area of crop?” (Citizen scientist participant, Year 1)*

Bloom and Crowder (2020)highlight the importance of understanding participants motivations for taking part in a citizen science project, based on their finding that this can predict the likelihood of a participant submitting data. At the end-point results event we asked participants to complete a final feedback survey (see Additional File 5). An online version of the survey was also distributed to all participants via email after the event. Eighty eight percent of our participants (n=17 respondents) reported that their motivation for taking part in the project was to learn about pollinators and all reported that this motivation was either moderately or very satisfied by the project. A similar proportion of respondents were also motivated ‘to help science’ and 93% reported this motivation was either moderately or very satisfied by participating in the project. Sixty four percent of respondents to the survey wanted to help protect the environment, and 58% wanted to track the food harvested and all these respondents reported their motivations to be either moderately or very satisfied. Other motivations for participating included being more observant, wanting to grow more crops and concerns regarding insect declines.

In terms of new information learnt about pollinators and food production since taking part in Team PollinATE, half of the participants that responded to the question (n=7) reported increased awareness of the importance and/or diversity of pollinators:

> *“I thought previously that bees were the only pollinators”* (Citizen science participant, Year 2)

as well as increased knowledge about which pollinators visit which crops (Additional File 5). One respondent also reported that they are more aware of their fellow allotment growers.

In the final survey we also asked participants to rate their ability to identify pollinators on a scale of 1 (not confident at all) to 10 (very confident), both before taking part in Team PollinATE and afterwards. The median rating was 3 before taking part in Team PollinATE and participants’ self-reported ratings increased significantly after taking part in the project (median = 7, W= 8, N=14, p<0.01, one-tailed). Had resources allowed, it would have been interesting to test this directly at the start and end of the project, and indeed this offers another potential option for data validation in citizen science projects (Ratnieks et al. 2016).

Twelve out of 14 respondents to the question “Have your gardening habits changed in any way since taking part in Team PollinATE?” responded positively, with the majority indicating that they now had increased awareness of pollinators in their allotments, and five respondents reported having adopted or expressed a desire to adopt insect-friendly behaviours, such as planting wildflowers and providing nesting habitat for pollinators:

> *“Much more aware and showing more interest in what insects are on what plants and a more ‘inclusive’ attitude to insects”* (Citizen science participant, Year 2)

One participant reported stopping using pesticides. Our findings add to the growing body of evidence suggesting that participation in citizen science projects can lead to both knowledge gain and behaviour change for participants. Given the huge potential for improving the sustainability of our cities (SDG11) and both urban and rural biodiversity (SDG15) through wildlife-friendly gardening and greenspace management as well as generating data for monitoring purposes citizen science has the potential to be used to both increase public awareness of and shift behaviour towards pro-environmental actions (Cosquer et al. 2012; Lewandowski and Oberhauser 2017; Griffiths-Lee et al. 2020; Sturm et al. 2021).

### Reflections and recommendations

Using land in urban areas to grow food has demonstrated benefits to wildlife such as insects, however little is currently known about how these insects may contribute to the pollination of flowering crops in urban farming, in part due to difficulties in accessing and monitoring these often *ad hoc* and dispersed habitats (Theodorou et al. 2017; Cohen et al. 2021). Here we evaluated the potential of using citizen science to enable urban growers themselves to collect data on the types of pollinating insect visiting their crops. Data collected by citizen scientists (CS) was compared against data collected by a trained researcher (R) using the same methodology, to assess the accuracy of data collected by non-specialists. We also collected feedback from participants throughout the project with the aim of improving data accuracy and the engagement and retention of participants, two of the major challenges associated with citizen science.

### Data accuracy

While the data on plant-pollinator interactions in urban areas collected by citizen scientists in this study could provide some indication of the insect groups most frequently observed (honeybees) and the most frequently visited crops (raspberries, squash) in a particular urban crop system, relying only on this dataset is inadvisable due to potential biases and inconsistencies in sampling between participants and inaccuracies in insect identification by non-specialists. In the current study this led to a more generalised network (i.e. crop plants receiving visits from more insect types) compared to data collected by a professional researcher. Participants also had concerns themselves regarding the usefulness and accuracy of their data, which they reported as a barrier to participation.

Here we validated the CS results by replicating the study with a professional researcher, which was time-consuming and could be difficult to execute over a larger geographical area and/or multiple urban cropping systems without a large research team. An alternative is to provide more in-depth or prolonged training in insect identification for participants, and perhaps also include an identification test that participants must pass before being permitted to submit data (Ratnieks et al. 2016). However, developing and delivering this training would still represent a substantial time investment for a single project lead or small project team. In-person training would not be possible for projects based across wider geographical areas, and developing a high-quality virtual resource could be time-consuming and costly. Taken together, this additional time investment could negate some of the potential cost-benefits from adopting a citizen science approach to data collection (Pocock et al. 2014; Encarnação et al. 2021; Parretti et al. 2023).

In-depth training also represents an additional time investment from participants which may be a barrier to involvement for some, given that lack of time was a major reason for not submitting data in the current study. However, having this additional pre-requirement may select for participants with greater capacity to commit to the project and the benefits vs. costs of this this will depend on whether consistency in participants and/or sampling sites over time is important for a particular project, or whether more *ad hoc* participation and data submission is acceptable. Other pollinator studies have required participants to submit photos for expert verification, but taking photos of fast-moving insects of high enough quality for identification can be challenging (Falk et al. 2019; Serret et al. 2022), requires access to technology and asking participants to upload photos can reduce the number of data submissions (Serret et al. 2022).

As a research team, we have had previous success with citizen science projects focussed on pollinating insects in urban contexts that did not rely on participants’ ability to identify insects, but which still provided important data regarding pollinator populations in urban food environments and the most appropriate management approaches (Birkin and Goulson 2015; Griffiths-Lee et al. 2020; Griffiths-Lee et al. 2022). For example, the UK-wide ‘Super Strawberries’ project (Griffiths-Lee et al. 2022), which arose from a question raised at the Year 1 feedback event. Team PollinATE participants raised concerns that they most frequently observed pollinating insects visiting ornamental flowers in their food growing spaces, and not crop flowers, and whether having these additional flowers might be ‘distracting’ insects from pollinating the crops. To test this, Super Strawberry participants were provided with two strawberry runners and seeds to grow the pollinator attractive, nectar rich plant borage (*Borago officinalis*). One strawberry plant was placed close to the borage, and one three meters away, and participants recorded the frequency of insect visits to the flowers of both, as well as the number and weights of fruit produced. This experiment was also repeated by a trained researcher and the results from the citizen scientist and researcher led experiment both concluded that planting strawberries alongside pollinator attractive flowers leads to better fruit yields. Since this study relied on participants only counting insects, and counting and weighing fruit, it may be a better approach for involving non-specialists in the study of pollination services in urban food growing.

### Participant engagement and retention

Aside from concerns regarding the accuracy and reliability of citizen scientist-collected data, another commonly reported drawback is the often-high levels of dropout of participants over the course of a project and the associated costs in terms of resource investment (Kleinke et al. 2018; Bloom and Crowder 2020). Only 18% of recruited participants submitted data in the current study, and only a handful submitted data consistently over the course of the two years. Protocol complexity and participant’s perceptions of method difficulty has previously been shown to affect the retention of volunteers and data submission rates in pollinator citizen science projects (Birkin and Goulson 2015; Bloom and Crowder 2020; Serret et al. 2022) but this can be countered by high levels of engagement, either in person or digitally, between participants and the project leads and/or between participants themselves, leading to a sense of community which can be a major motivation for some people for taking part in citizen science projects (Griffiths-Lee et al. 2022; Serret et al. 2022). The localised nature of TeamPollinATE, where participation was limited to residents of a single city, was beneficial in that it allowed us to collate in-person feedback regarding the experience of taking part in the project in the first year. Where possible we implemented the participants suggestions for improvement, for example by offering more hands-on training sessions at the start of the season, providing a message board on the project website where participants could provide each other with advice and support and post photos for identification, and providing more regular updates of collective findings across participants to counter the aforementioned feelings of decreased motivation when an individual failed to record any insects visiting crops in their individual plots.

Implementation of as many participant suggestions as possible led to 66% increase in data submissions between Year 1 and 2. Therefore we conclude that including a pilot stage and opportunities for participants to feedback on methods and barriers is key, and as others have suggested (Pocock et al. 2014; Bloom and Crowder 2020) surveying participants at the start of the recruitment process to understand their motivations for taking part could also be important, since this will assist with estimating how many participants should be recruited for adequate sample size, accounting for the likelihood of non-submissions (Domroese and Johnson 2017; Griffiths-Lee et al. 2023).

## Conclusion

In conclusion, this study highlights both the potential and limitations of working with growers to collect data on insect pollination in urban food systems. While the data collected by citizen scientists provided some insight into insect visitation patterns that overlapped with the researcher findings, such as the most frequently observed pollinator and most frequently visited crops, comparison of the data sets also revealed several biases, including overestimations of insect-crop generality and errors in species identification by citizen scientists. These discrepancies emphasize the importance of carefully validating citizen science data, especially when accurate species-level identification is required. Despite these challenges, the increase in data submissions after incorporating participant feedback demonstrates the importance of ongoing dialogue and flexibility in citizen science projects. Overall, citizen science can be a powerful tool for engaging the public in scientific research and increasing awareness and fostering behaviour change, such as improving awareness of pollinator diversity and more sustainable gardening practices. The inclusion of more focused training, simplification of data collection methods and improved support for participants may further enhance both the quality and quantity of data submitted, however, these require significant additional investment both from project organisers and participants.

## Acknowledgements

This work was funded by the Sussex Sustainability Research Programme and a UKRI Future Leaders Fellowship awarded to EN (MR/T021691/1). LS is supported by the UKRI funded UK Food Systems Centre for Doctoral Training studentship. AE was funded by a Royal Society Apex award. We are extremely grateful to all the citizen scientists who took part in the project and contributed to improving the pollinator survey methodology and training. We also thank the allotment site representatives and plot holders who provided site access for the researcher surveys. We would also like to thank the Brighton and Hove Allotment Federation (BHAF) and Brighton and Hove Organic Gardening Group (BHOGG) for their assistance with plot-holder recruitment. Thanks also to students Charlotte Cook and Emma Eatough and Jack McGee for assistance with the insect surveys, and Bryn Thomas of Brighton Permaculture Trust for assistance with tree identification.

## Author Contribution

EN, DG, PB, AE and HH designed the study and methodology. EN, DG, AE and HH collected data. EN, LS and SC analysed data and MCC provided advice and feedback on statistical analysis. EN wrote the first draft of the manuscript. All authors commented on the final draft of the manuscript.

## Data Availability Statement

The data that support the findings of this study are openly available in the supplementary materials of this paper.

## Research Ethics

Appropriate ethical approval for the study was obtained from the University of Sussex Cross Schools Research Ethics Committee (ER/EN97/1).

## References

Aizen, M.A., Garibaldi, L.A., Cunningham, S.A. and Klein, A.M. 2008. Long-term global trends in crop yield and production reveal no current pollination shortage but increasing pollinator dependency. Current Biology 18(20), pp. 1572–1575. doi: 10.1016/J.CUB.2008.08.066.

AJ Hamilton, K.B.H.-F.M.S.B.J.G.V.W. 2013. Give peas a chance? Urban agriculture in developing countries: a review. Agron Sustain Dev 34, pp. 45–73.

Altieri, M. 2009. Agroecology, small farms, and food sovereignty. Monthly Review 61, pp. 102–113.

Baldock, K. et al. 2019. A systems approach reveals urban pollinator hotspots and conservation opportunities. Nature Ecolgy & Evolution 3(3), p. 363.

Ballantyne, G., Baldock, K.C.R., Rendell, L. and Willmer, P.G. 2017. Pollinator importance networks illustrate the crucial value of bees in a highly speciose plant community. Scientific Reports 2017 7:1 7(1), pp. 1–13. Available at: https://www.nature.com/articles/s41598-017-08798-x [Accessed: 13 September 2022].

Lin, B.B.; P.S.M.; J.S. 2015. The future of urban agriculture and biodiversity-ecosystem services: challenges and next steps. Basic & Applied Ecology 16, pp. 189–201.

Bennett, A.B. and Lovell, S. 2019. Landscape and local site variables differentially influence pollinators and pollination services in urban agricultural sites. PLOS ONE 14(2), p. e0212034. Available at: https://journals.plos.org/plosone/article?id=10.1371/journal.pone.0212034 [Accessed: 27 February 2023].

Birkin, L. and Goulson, D. 2015. Using citizen science to monitor pollination services. Ecological Entomology 40(S1), pp. 3–11. Available at: https://onlinelibrary.wiley.com/doi/full/10.1111/een.12227 [Accessed: 15 August 2024].

Bloom, E.H. and Crowder, D.W. 2020. Promoting data collection in pollinator citizen science projects. Citizen Science: Theory and Practice 5(1). doi: 10.5334/CSTP.217.

Toomey, A.H.; D.M.C. 2013. Can citizen science lead to positive conservation. Human Ecology Review, pp. 50–62. Available at: https://www.proquest.com/docview/1511435864?sourcetype=Scholarly%20Journals [Accessed: 15 August 2024].

Cappa, F., Franco, S. and Rosso, F. 2022. Citizens and cities: Leveraging citizen science and big data for sustainable urban development. Business Strategy and the Environment 31(2), pp. 648–667. Available at: https://onlinelibrary.wiley.com/doi/full/10.1002/bse.2942 [Accessed: 14 August 2024].

Cohen, H., Philpott, S.M., Liere, H., Lin, B.B. and Jha, S. 2022. The relationship between pollinator community and pollination services is mediated by floral abundance in urban landscapes. Urban Ecosystems (24), pp. 275–290. Available at: 10.1007/s11252-020-01024-z [Accessed: 27 February 2023].

Colding, J., Lundberg, J. and Folke, C. 2006. Incorporating green-area user groups in urban ecosystem management. AMBIO:A journal of the human environment 35(5), pp. 237–244. Available at: https://bioone.org/journals/ambio-a-journal-of-the-human-environment/volume-35/issue-5/05-A-098R.1/Incorporating-Green-area-User-Groups-in-Urban-Ecosystem-Management/10.1579/05-A-098R.1.full [Accessed: 16 September 2022].

Cooley, H. and Vallejo-Marín, M. 2021. Buzz-Pollinated Crops: A Global Review and Meta-analysis of the Effects of Supplemental Bee Pollination in Tomato. Journal of Economic Entomology 114(2), pp. 505–519. Available at: 10.1093/jee/toab009 [Accessed: 15 August 2024].

Cosquer, A., Raymond, R. and Prevot-Julliard, A.C. 2012. Observations of Everyday Biodiversity: a New Perspective for Conservation? *Ecology and Society*, Published online: Oct 19, 2012 | doi:10.5751/ES-04955-170402 17(4). Available at: 10.5751/ES-04955-170402 [Accessed: 15 August 2024].

Dickinson, J.L. et al. 2012. The current state of citizen science as a tool for ecological research and public engagement. Frontiers in Ecology and the Environment 10(6), pp. 291–297. Available at: https://onlinelibrary.wiley.com/doi/full/10.1890/110236 [Accessed: 15 August 2024].

Dobson, M.C., Crispo, M., Blevins, R.S., Warren, P.H. and Edmondson, J.L. 2021a. An assessment of urban horticultural soil quality in the United Kingdom and its contribution to carbon storage. Science of The Total Environment 777, p. 146199. doi: 10.1016/J.SCITOTENV.2021.146199.

Dobson, M.C., Reynolds, C., Warren, P.H. and Edmondson, J.L. 2021b. “My little piece of the planet”: the multiplicity of well-being benefits from allotment gardening. British Food Journal 123(3), pp. 1012–1023. doi: 10.1108/BFJ-07-2020-0593/FULL/XML.

Domroese, M.C. and Johnson, E.A. 2017. Why watch bees? Motivations of citizen science volunteers in the Great Pollinator Project. Biological Conservation 208, pp. 40–47. doi: 10.1016/J.BIOCON.2016.08.020.

Druschke, C.G. and Seltze, C.E. 2012. Failures of Engagement: Lessons Learned from a Citizen Science Pilot Study. Applied Environmental Education & Communication 11(3–4), pp. 178–188. Available at: https://www.tandfonline.com/doi/abs/10.1080/1533015X.2012.777224 [Accessed: 15 August 2024].

Dunkley, R.A. 2017. The Role of Citizen Science in Environmental Education: A Critical Exploration of the Environmental Citizen Science Experience. In: AnalyzingThe Role Of Citizen Science In Modern Research. IGI Global, pp. 213–230. Available at: https://www.igi-global.com/chapter/the-role-of-citizen-science-in-environmental-education/170191 [Accessed: 15 August 2024].

Edmondson, J.L. et al. 2020. The hidden potential of urban horticulture. Nature Food 2020 1:3 1(3), pp. 155–159. Available at: https://www.nature.com/articles/s43016-020-0045-6 [Accessed: 13 September 2022].

Encarnação, J., Baptista, V., Teodósio, M.A. and Morais, P. 2021. Low-Cost Citizen Science Effectively Monitors the Rapid Expansion of a Marine Invasive Species. Frontiers in Environmental Science 9, p. 752705. Available at: www.frontiersin.org [Accessed: 14 August 2024].

F., H. 2018. DHARMa: Residual Diagnostics for Hierarchical (Multi-Level / Mixed) Regression Models. R Packag version 020. Available at: https://cir.nii.ac.jp/crid/1370580229833186830.bib?lang=en [Accessed: 15 August 2024].

Falk, S. et al. 2019. Evaluating the ability of citizen scientists to identify bumblebee (Bombus) species. PLOS ONE 14(6), p. e0218614. Available at: https://journals.plos.org/plosone/article?id=10.1371/journal.pone.0218614 [Accessed: 14 August 2024].

Fraisl, D. et al. 2020. Mapping citizen science contributions to the UN sustainable development goals. Sustainability Science 15(6), pp. 1735–1751. Available at: https://link.springer.com/article/10.1007/s11625-020-00833-7 [Accessed: 14 August 2024].

Fritz, S. et al. 2019. Citizen science and the United Nations Sustainable Development Goals. Nature Sustainability 2019 2:10 2(10), pp. 922–930. Available at: https://www.nature.com/articles/s41893-019-0390-3 [Accessed: 14 August 2024].

Fründ, J., Mccann, K.S. and Williams, N.M. 2016. Sampling bias is a challenge for quantifying specialization and network structure: lessons from a quantitative niche model. Oikos 125(4), pp. 502–513. Available at: https://onlinelibrary.wiley.com/doi/full/10.1111/oik.02256 [Accessed: 27 February 2023].

Garratt, M.P.D., Potts, S.G., Banks, G., Hawes, C., Breeze, T.D., O’Connor, R.S. and Carvell, C. 2019a. Capacity and willingness of farmers and citizen scientists to monitor crop pollinators and pollination services. Global Ecology and Conservation 20, p. e00781. doi: 10.1016/J.GECCO.2019.E00781.

Garratt, M.P.D., Potts, S.G., Banks, G., Hawes, C., Breeze, T.D., O’Connor, R.S. and Carvell, C. 2019b. Capacity and willingness of farmers and citizen scientists to monitor crop pollinators and pollination services. Global Ecology and Conservation 20, p. e00781. doi: 10.1016/J.GECCO.2019.E00781.

Griffiths-Lee, J., Nicholls, E. and Goulson, D. 2020. Companion planting to attract pollinators increases the yield and quality of strawberry fruit in gardens and allotments. Ecological Entomology 45(5), pp. 1025–1034. Available at: https://onlinelibrary.wiley.com/doi/full/10.1111/een.12880 [Accessed: 20 September 2022].

Griffiths-Lee, J., Nicholls, E. and Goulson, D. 2022. Sown mini-meadows increase pollinator diversity in gardens. Journal of Insect Conservation 26(2), pp. 299–314. Available at: https://link.springer.com/article/10.1007/s10841-022-00387-2 [Accessed: 15 August 2024].

Griffiths-Lee, J., Nicholls, E. and Goulson, D. 2023. Sow Wild! Effective Methods and Identification Bias in Pollinator-Focused Experimental Citizen Science. Citizen Science: Theory and Practice 8(1). doi: 10.5334/CSTP.550.

Holligan, B. and Howe, H. 2024. How property relations shape experiences and transformative potential of urban growing spaces: Connecting land, food, and Earth justice perspectives. Elementa 12(1). Available at: /elementa/article/12/1/00082/202563/How-property-relations-shape-experiences-and [Accessed: 14 August 2024].

Borysiak, J., Mizgajski, A. and Speak, A. 2017. Floral biodiversity of allotment gardens and its contribution to urban green infrastructure. Urban Ecosystems 20, pp. 323–335.

Quesada, J. and MacGregor–Fors, I. 2010. Avian community responses to the establishment of small garden allotments within a Mediterranean habitat mosaic. Animal Biodiversity and Conservation 33, pp. 53–61.

Kleinke, B., Prajzner, S., Gordon, C., Hoekstra, N., Kautz, A. and Gardiner, M. 2018. Identifying Barriers to Citizen Scientist Retention When Measuring Pollination Services. Citizen Science: Theory and Practice 3(1), pp. 2–2. Available at: https://account.theoryandpractice.citizenscienceassociation.org/index.php/up-j-cstp/article/view/cstp.99 [Accessed: 14 August 2024].

Koffler, S., Barbiéri, C., Ghilardi-Lopes, N.P., Leocadio, J.N., Albertini, B., Francoy, T.M. and Saraiva, A.M. 2021. A Buzz for Sustainability and Conservation: The Growing Potential of Citizen Science Studies on Bees. Sustainability 2021, Vol. 13, Page 959 13(2), p. 959. Available at: https://www.mdpi.com/2071-1050/13/2/959/htm [Accessed: 14 August 2024].

Kosmala, M., Wiggins, A., Swanson, A. and Simmons, B. 2016. Assessing data quality in citizen science. Frontiers in Ecology and the Environment 14(10), pp. 551–560. Available at: https://onlinelibrary.wiley.com/doi/full/10.1002/fee.1436 [Accessed: 15 August 2024].

Kremen, C., Ullman, K.S. and Thorp, R.W. 2011. Evaluating the Quality of Citizen-Scientist Data on Pollinator CommunitiesEvaluación de la Calidad de Datos de Comunidades de Polinizadores Tomados por Ciudadanos-Científicos. Conservation Biology 25(3), pp. 607–617. Available at: https://onlinelibrary.wiley.com/doi/full/10.1111/j.1523-1739.2011.01657.x [Accessed: 14 August 2024].

Laughton, R. 2017. A Matter of Scale: A study of the productivity, financial viability and multifunctional benefits of small farms. *Centre for Agroecology, Water and Resilience (CAWR)*, Coventry.

Levin, M. and Kuehl, R. 1968. Comparison of three sampling methods for estimating honey bee visitation to flowers of cucumbers. Journal of Economic Entomology (67), pp. 1487–1489. Available at: https://academic.oup.com/jee/article-abstract/61/6/1487/972512 [Accessed: 27 February 2023].

Lewandowski, E.J. and Oberhauser, K.S. 2017. Butterfly citizen scientists in the United States increase their engagement in conservation. Biological Conservation 208, pp. 106–112. doi: 10.1016/J.BIOCON.2015.07.029.

Lin Hunter, D.E., Newman, G.J. and Balgopal, M.M. 2023. What’s in a name? The paradox of citizen science and community science. Frontiers in Ecology and the Environment 21(5), pp. 244–250. Available at: https://onlinelibrary.wiley.com/doi/full/10.1002/fee.2635 [Accessed: 14 August 2024].

Lukyanenko, R., Wiggins, A. and Rosser, H.K. 2020. Citizen Science: An Information Quality Research Frontier. Information Systems Frontiers 22(4), pp. 961–983. Available at: https://link.springer.com/article/10.1007/s10796-019-09915-z [Accessed: 15 August 2024].

MacPhail, V.J. and Colla, S.R. 2020. Power of the people: A review of citizen science programs for conservation. Biological Conservation 249, p. 108739. doi: 10.1016/J.BIOCON.2020.108739.

Maher, S., Manco, F. and Ings, T.C. 2019. Using citizen science to examine the nesting ecology of ground-nesting bees. Ecosphere 10(10), p. e02911. Available at: https://onlinelibrary.wiley.com/doi/full/10.1002/ecs2.2911 [Accessed: 15 August 2024].

Mason, L. and Arathi, H.S. 2019. Assessing the efficacy of citizen scientists monitoring native bees in urban areas. Global Ecology and Conservation 17, p. e00561. doi: 10.1016/J.GECCO.2019.E00561.

Matthies, S., Rüter, S., Prasse, F. and Schaarschmidt F. 2015. Factors driving the vascular plant species richness in urban green spaces: Using a multivariable approach. Landscape Urban Planning 134, pp. 177–187.

McDougall, R., Kristiansen, P. and Rader, R. 2019. Small-scale urban agriculture results in high yields but requires judicious management of inputs to achieve sustainability. Proceedings of the National Academy of Sciences of the United States of America 116(1), pp. 129–134. Available at: https://www.pnas.org/doi/abs/10.1073/pnas.1809707115 [Accessed: 13 September 2022].

Nicholls, E., Ely, A., Birkin, L., Basu, P. and Goulson, D. 2020. The contribution of small-scale food production in urban areas to the sustainable development goals: a review and case study. Sustainability Science 15(6), pp. 1585–1599. Available at: https://link.springer.com/article/10.1007/s11625-020-00792-z [Accessed: 16 September 2022].

Nicholls, E., Griffiths-Lee, J., Basu, P., Chatterjee, S. and Goulson, D. 2023. Crop–pollinator interactions in urban and peri-urban farms in the United Kingdom. Plants People Planet 5(5). doi: 10.1002/ppp3.10376.

Oxford English Dictionary. [no date]. Available at: https://www.oed.com/?tl=true [Accessed: 15 August 2024].

Parretti, P. et al. 2023. Citizen Science and Expert Judgement: A Cost-Efficient Combination to Monitor and Assess the Invasiveness of Non-Indigenous Fish Escapees. Journal of Marine Science and Engineering 11(2), p. 438. Available at: https://www.mdpi.com/2077-1312/11/2/438/htm [Accessed: 14 August 2024].

Peter, M., Diekötter, T., Höffler, T. and Kremer, K. 2021. Biodiversity citizen science: Outcomes for the participating citizens. People and Nature 3(2), pp. 294–311. Available at: https://onlinelibrary.wiley.com/doi/full/10.1002/pan3.10193 [Accessed: 15 August 2024].

Pocock, M.J.O., Chapman, D.S., Sheppard, L.J. and Roy, H.E. 2014. Choosing and using citizen science: a guide to when and how to use citizen science to monitor biodiversity and the environment. NERC/Centre for Ecology & Hydrology.

Rader, R. et al. 2016. Non-bee insects are important contributors to global crop pollination. Proceedings of the National Academy of Sciences of the United States of America 113(1), pp. 146–151. Available at: https://www.pnas.org/doi/abs/10.1073/pnas.1517092112 [Accessed: 15 August 2024].

Ratnieks, F.L.W., Schrell, F., Sheppard, R.C., Brown, E., Bristow, O.E. and Garbuzov, M. 2016. Data reliability in citizen science: learning curve and the effects of training method, volunteer background and experience on identification accuracy of insects visiting ivy flowers. Methods in Ecology and Evolution 7(10), pp. 1226–1235. Available at: https://onlinelibrary.wiley.com/doi/full/10.1111/2041-210X.12581 [Accessed: 15 August 2024].

Serret, H., Deguines, N., Jang, Y., Lois, G. and Julliard, R. 2019. Data quality and participant engagement in citizen science: comparing two approaches for monitoring pollinators in France and South Korea. Citizen Science: Theory and Practice (4), p. 22. Available at: https://hal.science/hal-02553137/document [Accessed: 14 August 2024].

Sturm, U., Straka, T.M., Moormann, A. and Egerer, M. 2021. Fascination and Joy: Emotions Predict Urban Gardeners’ Pro-Pollinator Behaviour. Insects 2021, Vol. 12, Page 785 12(9), p. 785. Available at: https://www.mdpi.com/2075-4450/12/9/785/htm [Accessed: 15 August 2024].

Thebo, A.L., Drechsel, P. and Lambin, E.F. 2014. Global assessment of urban and peri-urban agriculture: irrigated and rainfed croplands. Environmental Research Letters 9(11), p. 114002. Available at: https://iopscience.iop.org/article/10.1088/1748-9326/9/11/114002 [Accessed: 13 September 2022].

Theodorou, P., Albig, K., Radzevičiūtė, R., Settele, J., Schweiger, O., Murray, T.E. and Paxton, R.J. 2017. The structure of flower visitor networks in relation to pollination across an agricultural to urban gradient. Functional Ecology 31(4), pp. 838–847. Available at: https://onlinelibrary.wiley.com/doi/full/10.1111/1365-2435.12803 [Accessed: 27 February 2023].

Wals, A.E.J., Brody, M., Dillon, J. and Stevenson, R.B. 2014. Convergence between science and environmental education. Science 344(6184), pp. 583–584. Available at: https://www.science.org/doi/10.1126/science.1250515 [Accessed: 15 August 2024].

Zezza, A. and Tasciotti, L. 2008. Does Urban Agriculture Enhance Dietary Diversity? Empirical Evidence from a Sample of Developing Countries. Available at: https://ageconsearch.umn.edu/record/44390 [Accessed: 27 February 2023].

