## Supplemental Materials for "Understanding pollination in urban food production: the importance of data validation and participant feedback for citizen science project design"

***Plants, People, Planet* Supporting Information**

**The following Supporting Information is available for this article:**

**Table S1** Total number of plots and observations made by citizen scientists and researchers at each allotment site or individual garden.

**Table S2** Results of the sticker voting at the Year 1 mid-point feedback event.

**Table S3** Comments left by project participants on the comment boards at the Year 1 mid-point feedback event and grouped by key themes identified through thematic analysis.

**Table S4** Key topics and themes that emerged from the Year 1 mid-point feedback event World Café with project participants.

**Table S5** Total number of observations made by researchers and citizen scientists of each crop type, the mean number of plants of each crop surveyed per twice monthly observation, the mean number of flowers per crop.

**Table S6-** Z-test with Bonferroni corrections

**Table S7** Total number of observations made by researchers and citizen scientists per month

**Table S9** Results of glm

**Table S10** Results of post-hoc comparisons

**Table S11** Z-test on proportion of observations of each insect group

**Table S12** Species indicator analysis

**Table S13** Species and insect group-level metrics for each visitation network.

**Methods S1** Survey instructions for participants

**Methods S2** Plant-pollinator interactions analysis

**Code S1** Code used to analyse plant-pollinator data (R Script submitted separately)

**Dataset S1** Participants responses to endpoint feedback survey (Excel file submitted separately)

**Dataset S2** Pollinator observations by citizen scientists and researchers (Excel file submitted separately)

**Additional File S1-** Welcome pack for participants (PDF submitted separately)

**Table S1** Total number of plots and observations made by citizen scientists and researchers at each allotment site or individual garden.

| **Allotment Site** | **Plot Type** | **N Citizen Scientist plots** | **N Crop Obs.** | **N Researcher Plots** | **N Crop Obs.** |
|---|---|---|---|---|---|
| Camp Site | Individual | 2 | 29 | 1 | 2 |
| Coldean | Individual | 1 | 39 | 0 | 0 |
| Eastbrook Farm | Individual | 1 | 3 | 0 | 0 |
| Hoggs Platt | Individual | 1 | 16 | 1 | 35 |
| Keston | Individual | 1 | 22 | 18 | 86 |
| Manton Road | Individual | 1 | 7 | 0 | 0 |
| Moulsecoomb Forest Garden | Community | 1 | 14 | 1 | 73 |
| Moulsecoomb Estate | Individual | 2 | 89 | 1 | 5 |
| Racehill Community Orchard | Community | 1 | 3 | 1 | 7 |
| Roedale Valley | Individual | 4 | 116 | 29 | 304 |
| Stanford Community Gardens | Community | 1 | 21 | 0 | 0 |
| Tenantry Down | Individual | 1 | 6 | 0 | 0 |
| Upper Roedale | Individual | 2 | 53 | 0 | 0 |
| Weald | Individual | 7 | 79 | 18 | 390 |
| Whitehawk | Individual | 1 | 4 | 13 | 222 |
| Gardens | Individual | 6 | 39 | 0 | 0 |
| | **Total** | **33** | **540** | **83** | **1124** |

**Table S2** Results of the sticker voting at the Year 1 mid-point feedback event.

| Sticker Vote Question | Participant Responses | | | |
|---|---|---|---|---|
| 1. How would you rate the level of communication from the Team PollinATE organisers? | | ***Too Little*** | ***Just Right*** | ***Too much*** |
| | N | 0 | 8 | 0 |
| | % | 0% | 100% | 0% |
| 2. What was your favourite part of Team PollinATE? | | ***Pollinator surveys*** | ***Yield tracking*** | ***Pest Control Diaries*** |
| | N | 6 | 2 | 0 |
| | % | 75% | 25% | 0% |
| 3. What was your least favourite part of Team PollinATE? | N | 1 | 1 | 3 |
| | % | 20% | 20% | 60% |
| 4. How easy were the data collection methods? | | ***Very Easy*** | ***Moderately Easy*** | ***Moderately Hard*** |
| | N | 5 | 3 | 1 |
| | % | 56% | 33% | 11% |
| 5. How easy was it to submit data? | N | 3 | 2 | 2 |
| | % | 42% | 29% | 29% |

**Table S3** Participant feedback at the Year 1 mid-point feedback event, collected via anonymous written comments left under the four key question headings. The comments have been grouped according to thematic analysis.

| | |
|---|---|
| *'It made me think even more about pollinators and look more closely at the insects on my plot'* | **Awareness** |
| *'Learning about pollinators and becoming more observant'* | Awareness |
| *'Looking closely at pollinating insects- trying to identify different bees'* | Awareness |
| *'It made me more aware of the insects that pollinate our crops'* | Awareness |
| *'Makes me think about bees more and watch them closer'* | Awareness |
| *'Yield monitor'* | Data collection |
| *'It was great finding out that we predominantly have honeybees'* | Knowledge |
| *'Learning about pollinators'* | Knowledge |
| *'Being part of an important piece of research'* | Participation |
| *'Enthusiasm of team'* | Participation |
| **Q2. What didn't you like about taking part in Team PollinATE?** | **Key Themes** |
| *'Had planted bee loving flowers and bees more on flowers so felt that skewed the data'* | Data accuracy concerns |
| *'Found counting blossoms and insects very difficult on densely planted plot'* | Methods too difficult |
| *'Found counting number of plants difficult to do meaningfully'* | Methods too difficult |
| *'Reporting survey results and crop yields was fiddly- would be good to have individual account to save entering [personal] details each time'* | Support for data collection and submission |
| *'Maybe I'm very stupid with computers but each time I had to start at the beginning again with site location questions'* | Support for data collection and submission |
| **Q3. What made it difficult to collect data?** | **Key Themes** |
| *'Bothered me how inaccurate the data was- e.g. impossible to count raspberry flowers'* | Data accuracy concerns |
| *'Weather'* | Environmental effects |
| *'The bees were mostly on the non-crop flowers/plants- difficult NOT counting them'* | Lack of data |
| *'Collecting data at times when trying to keep everything alive and pushed for time'* | Lack of time |
| *'Counting flowers- inputting results when I want to avoid the computer'* | Methods too difficult |
| *'Counting plants with multiple blooms'* | Methods too difficult |
| *'Counting the flowers, esp. on crops with a mass of flowers'* | Methods too difficult |
| *'Knowing whether to count bees that were utilising 'robber-bee' entry holes (most of them at the runner beans)* | Support for data collection and submission |
| *'Time! Or rather prioritising data collection over weeding, watering, planting etc.'* | Lack of time |
| **Q4. How could we make Team PollinATE better for next year?** | **Key Themes** |
| *'Collect info about use/not use of pesticides (or did you??)'* | Improved communication |
| *'Better informed'* | Improved communication |
| *'Yield monitor- an organic equivalent '* | Methodological suggestion |
| *'Separate onions with spring onions'* | Methodological suggestion |
| *'Possibly a more flexible survey schedule- depressing when got behind!'* | Participant motivation and retention |
| *Having a 'log in' so no need to repeat some entry info each time* | Support for data collection and submission |
| *'Identity online log in'* | Support for data collection and submission |
| *'Online forum for discussion'* | Support for data collection and submission |

**Table S4** Key topics and themes that emerged from thematic analysis of the World Café conducted at mid-point feedback event. Participants circulated between two tables, with two broad questions 1) What is your experience of being a Team PollinATE volunteer and 2) What do you think of the questions being investigated by Team PollinATE.

| **World café Table 1: What is your experience of being Team PollinATE volunteers (led by Prof. Dave Goulson)** | |
|---|---|
| **Subquestion 1) What were/are your motivations for taking part in Team PollinATE?** | **Key Themes** |
| *Gathering data and excitement about doing citizen science* | Data collection |
| *Enjoyed collecting data- moment to step back and admire their plot* | Data collection |
| *Concern over decline of bees and industrial food production* | Environmental concerns |
| *Love of bees* | Interest in topic |
| *Interest in pollinators and wanted to learn about more than just honeybees and bumblebees* | Interest in topic |
| *Interest in pest monitoring component and how much food can be grown the natural way* | Interest in topic |
| *Curiosity* | Interest in topic |
| **Subquestion 2) What were the barriers to you taking part/submitting data?** | **Key Themes** |
| *Concerns about missing a survey and data not being useful* | Data accuracy concerns |
| *Not counting insects was very discouraging* | Lack of data |
| *Often no bees* | Lack of data |
| *Too busy- life takes over* | Lack of time |
| *Process very demanding and time is already limited at the allotment and long list of things to do* | Lack of time |
| *Too many crops* | Methods too difficult |
| *Hard- couldn't estimate flower numbers and felt data would be far too inaccurate* | Methods too difficult |
| *Initial effort to learn* | Methods too difficult |
| *Struggled with lack of efficiency in entering information* | Methods too difficult |
| *Going on computer off-putting* | Support for data collection and submission |
| *No scales for heavy crops* | Support for data collection and submission |
| **Subquestion 3) Do you have any suggestions for improvements to Team PollinATE for following years?** | **Key Themes** |
| *Herbs are very high value, include in yield calculator?* | Methodological suggestion |
| *Photos of crops for yields* | Methodological suggestion |
| *Why not start in January? some early crops e.g. gooseberries and pollinators are active* | Methodological suggestion |
| *Weigh veg when cleaning up* | Methodological suggestion |
| *People may like to submit data in different ways- paper vs. laptop* | Methodological suggestion |
| *Board at entrance to put up results, or highlight interesting observations* | Participant motivation |
| *Monthly update to keep up enthusiasm* | Participant motivation |
| *Pressure to keep up with data collection- make clear it doesn't matter if miss a week* | Participant motivation |
| *Posters at allotments* | Recruitment |
| *Recruitment- use allotment email lists* | Recruitment |
| *Results online to download and share with people- add to flyer so people can see what they would be contributing towards* | Recruitment |
| *Advice on estimating flower numbers and reassurance that it won't be highly accurate* | Support for data collection and submission |
| *Ambassadors would be great* | Support for data collection and submission |
| *Is there somewhere to upload photos so we can check we're doing it right e.g. community forum?* | Support for data collection and submission |
| *More help on insect identification* | Support for data collection and submission |
| *Upload button hard to find on website* | Support for data collection and submission |

| World Café Table 2: What do you think of the questions being investigated by Team PollinATE (led by Dr Beth Nicholls) | |
|---|---|
| **Subquestion 1) Do you have any general comments on the aims of Team PollinATE?** | **Key Themes** |
| *Concerns over accuracy/meaningfulness of the data* | Data accuracy concerns |
| *Data is a good baseline/starting point for a first year* | Long term data |
| *Long term data is very important if we are going to observe changes/problems/cycles in pest populations* | Long term data |
| *Hard to measure size of plot- additional guidance needed* | Methods too difficult |
| *Providing kit such as weighing scales would be useful* | Methodological suggestion |
| **Subquestion 2) Comments on Aim 1) Understanding which insects pollinate crops grown in urban areas** | **Key Themes** |
| *Nocturnal pollinators* | Interest in topic |
| *What is the effect of growing bee friendly flowers on crop visitation by pollinators* | Interest in topic |
| *Would like some recommendations of what to grow/do to help pollinators and wildlife* | Interest in topic |
| *Curious about relationship between pollinations and yield since not seeing many pollinators on crop* | Interest in topic |
| *Quantify area 'cultivated' (allotment rules- does 75% include recreation/flowers or not?)- particularly true if opening project to gardeners.* | Methodological suggestion |
| **Subquestion 3) Comments on Aim 2) Quantifying how much food can be grown in urban areas** | **Key Themes** |
| *Often grow more than I harvest- 'yields' may not be accurate* | Data accuracy concerns |
| *What proportion of total diet is food grown on allotment?* | Diet |
| *Does allotment increase diet diversity- grow things that you can't buy in shops. Growing my own has made my diet more seasonal (like when we were kids).* | Diet |
| *Environmental effects on yields, as well as pollination- e.g. frost/rain/location/soil type* | Interest in topic |
| *Rather than weighing, use volume e.g. number of punnets/bags* | Methodological suggestion |
| *Herbs, cut flowers also grown to save money* | Methodological suggestion |
| *Would it be possible to assign people different crops rather than measuring from all to reduce burden* | Methodological suggestion |
| *Should we just log edible fruit or should there be a differentiation?* | Methodological suggestion |
| *Could we use phone photos to estimate yield?* | Methodological suggestion |
| *The bigger my yields- less inclined to record* | Participant motivation |
| **Subquestion 4) Comments on Aim 3) Determining which are the most common pests in urban areas and how are they controlled** | **Key Themes** |
| *Might some people feel 'bad' about pesticide use and not be honest?* | Data accuracy concerns |
| *What about predators of pollinators (pests)- compare with bird monitoring by rangers/biosphere officers* | Interest in topic |
| *How do allotment pests vary by site location e.g. soil type, proximity to sea or woodland* | Interest in topic |
| *How far do pests travel between plots?* | Methodological suggestion |
| *Amount of pesticides used vs. effectiveness* | Methodological suggestion |
| *Are you making a value judgement about pesticide use?* | Researcher motivations |

**Table S5** Total number of observations made by researchers and citizen scientists of each crop type, the mean number of plants of each crop surveyed per twice monthly observation, the mean number of flowers per crop and the mean number of insects observed per crop flower.

| Observer Type | Crop | Surveys | Mean ± SD Plants | Mean ± SD Flowers | Mean ± SD Insects | Mean ± SD Insects/Flower |
|---|---|---|---|---|---|---|
| **Citizen Scientists** | Apple | 26 | 2.46± 1.70 | 427.08± 511.26 | 1.96±2.27 | 0.02±0.07 |
| | Broad bean | 53 | 26.09±26.16 | 311.47±437.06 | 1.57±1.91 | 0.03±0.06 |
| | Raspberry | 94 | 15.29±28.01 | 134.61±270.44 | 5.30±5.50 | 0.18±0.33 |
| | Cherry | 17 | 2.29±1.26 | 355.41±343.72 | 0.82±1.63 | 0.00±0.01 |
| | Squash | 128 | 3.90±3.75 | 9.61±11.18 | 2.90±5.26 | 0.72±2.30 |
| | Cucumber | 43 | 3.19±2.08 | 30.16±31.62 | 1.07±1.80 | 0.04±0.08 |
| | Currants | 28 | 5.14±4.22 | 647.14±915.52 | 1.64±2.30 | 0.01±0.02 |
| | Runner bean | 75 | 12.64±8.81 | 170.56±225.48 | 3.61±5.96 | 0.05±0.10 |
| | Strawberry | 69 | 18.06±15.92 | 148.34±207.43 | 1.77±2.82 | 0.14±0.84 |
| | Tomato | 60 | 8.40±8.22 | 63.18±74.80 | 0.83±1.36 | 0.04±0.14 |
| **Researcher** | Apple | 133 | 1.83±2.03 | 188.29±288.30 | 1.09±1.92 | 0.01±0.02 |
| | Broad bean | 94 | 21.27±21.52 | 139.43±169.58 | 0.37±0.72 | 0.01±0.04 |
| | Raspberry | 326 | 28.20±34.24 | 120.66±167.93 | 2.60±3.31 | 0.04±0.06 |
| | Cherry | 50 | 1.22±0.59 | 292.26±461.54 | 1.42±2.03 | 0.02±0.03 |
| | Squash | 375 | 3.08±3.38 | 5.06±13.54 | 1.15±2.60 | 0.36±1.06 |
| | Cucumber | 75 | 2.84±1.81 | 9.75±12.32 | 0.83±3.77 | 0.12±0.61 |
| | Currants | 79 | 3.13±2.56 | 324.05±266.61 | 0.65±1.12 | 0.00±0.00 |
| | Runner bean | 236 | 8.98±8.03 | 64.40±72.18 | 0.94±1.51 | 0.02±0.04 |
| | Strawberry | 124 | 11.70±14.51 | 43.34±65.62 | 0.44±0.94 | 0.04±0.16 |
| | Tomato | 87 | 6.84±7.71 | 40.76±51.85 | 0.06±0.23 | 0.00±0.01 |

**Table S6** Z-test (with Bonferroni corrections) comparing the proportion of total observations of each crop type between citizen scientist and researcher dataset.

| Crop | Z_value | P_value | Adjusted_P_value |
|---|---|---|---|
| Apple | -3.219 | 0.001 | 0.013 |
| Broad bean | 2.467 | 0.014 | 0.136 |
| Raspberry | -2.521 | 0.012 | 0.117 |
| Cherry | -0.360 | 0.719 | 1.000 |
| Squash | -1.065 | 0.287 | 1.000 |
| Cucumber | 2.291 | 0.022 | 0.219 |
| Currants | -0.270 | 0.787 | 1.000 |
| Runner bean | -1.363 | 0.173 | 1.000 |
| Strawberry | 2.760 | 0.006 | 0.058 |
| Tomato | 3.809 | 0.000 | 0.001 |

**Table S7** Total and mean number of surveys, and the mean number of plants, flowers and insects observed by researchers and citizen scientists per month

| Observer Type | Month | Surveys | Mean ±SD Plants | Mean ±SD Flowers | Mean ±SD Insects | Mean±SD Insects/Flower |
|---|---|---|---|---|---|---|
| **Citizen Scientist** | April | 42 | 13.33 ± 20.47 | 476.31±793.31 | 1.10±1.96 | 0.02±0.06 |
| | May | 121 | 15.70±20.50 | 320.90±411.20 | 2.00±3.20 | 0.03±0.10 |
| | June | 148 | 10.35±15.03 | 97.33±178.83 | 2.75±4.16 | 0.24±0.60 |
| | July | 135 | 8.48±14.79 | 95.54±223.83 | 2.74±3.52 | 0.20±0.61 |
| | August | 104 | 9.40±17.33 | 46.38±69.36 | 3.24±5.50 | 0.30±0.81 |
| | September | 30 | 7.10±10.01 | 67.93±190.87 | 3.53±8.99 | 0.94±4.45 |
| **Researcher** | April | 101 | 4.55±9.28 | 245.46±352.93 | 0.95±1.74 | 0.01±0.03 |
| | May | 357 | 9.28±14.87 | 171.87±255.60 | 1.11±2.32 | 0.01±0.04 |
| | June | 240 | 11.95±19.45 | 86.56±150.57 | 1.44±2.75 | 0.21±1.05 |
| | July | 283 | 10.82±21.32 | 41.18±72.30 | 1.54±3.46 | 0.22±0.08 |
| | August | 513 | 11.65±21.19 | 43.68±73.39 | 1.14±1.92 | 0.08±0.22 |
| | September | 85 | 13.04±27.03 | 40.71±90.95 | 0.78±1.52 | 0.08±0.21 |

**Table S8** Z-test (with Bonferroni corrections) comparing the proportion of total observations conducted each month between citizen scientist and researcher dataset.

| Crop | Z_value | P_value | Adjusted_P_value |
|---|---|---|---|
| April | 0.700 | 0.484 | 1.000 |
| May | -0.867 | 0.386 | 1.000 |
| June | 5.535 | 0.000 | 0.000 |
| July | 2.790 | 0.005 | 0.032 |
| August | -6.637 | 0.000 | 0.000 |
| September | -0.193 | 0.847 | 1.000 |

**Table S9** Comparison between the researcher and citizen science datasets in terms of the number of insect visits observed across sampling months and crops. We conducted a glm with a negative binomial distribution, to handle overdispersion in the data, using the formula (Total Insects Observed ~ Observer Type * Crop Type + Observer Type * Sampling Month).

| term | estimate | std.error | statistic | p.value |
|---|---|---|---|---|
| (Intercept) | 0.384 | 0.373 | 1.029 | 0.303 |
| Researcher | -0.360 | 0.457 | -0.787 | 0.431 |
| Broad Bean | -0.460 | 0.368 | -1.249 | 0.212 |
| Raspberry | 0.626 | 0.357 | 1.754 | 0.080 |
| Cherry | -0.720 | 0.517 | -1.391 | 0.164 |
| Squash | -0.135 | 0.373 | -0.361 | 0.718 |
| Cucumber | -1.117 | 0.426 | -2.619 | **0.009** |
| Currants | -0.098 | 0.407 | -0.240 | 0.810 |
| Runner bean | 0.061 | 0.387 | 0.158 | 0.874 |
| Strawberry | -0.251 | 0.353 | -0.711 | 0.477 |
| Tomato | -1.304 | 0.402 | -3.242 | **0.001** |
| May | 0.363 | 0.301 | 1.205 | 0.228 |
| June | 0.605 | 0.336 | 1.799 | 0.072 |
| July | 0.810 | 0.349 | 2.319 | **0.020** |
| August | 0.911 | 0.358 | 2.545 | **0.011** |
| September | 0.946 | 0.428 | 2.211 | **0.027** |
| Researcher: Broad Bean | -0.646 | 0.454 | -1.423 | 0.155 |
| Researcher: Raspberry | 0.268 | 0.415 | 0.646 | 0.518 |
| Researcher: Cherry | 1.035 | 0.602 | 1.718 | 0.086 |
| Researcher: Squash | 0.170 | 0.441 | 0.385 | 0.700 |
| Researcher: Cucumber | 0.661 | 0.519 | 1.275 | 0.202 |
| Researcher: Currants | -0.399 | 0.483 | -0.826 | 0.409 |
| Researcher: Runner bean | -0.153 | 0.458 | -0.333 | 0.739 |
| Researcher: Strawberry | -0.626 | 0.422 | -1.484 | 0.138 |
| Researcher: Tomato | -1.718 | 0.652 | -2.636 | **0.008** |
| Researcher: May | -0.293 | 0.388 | -0.756 | 0.450 |
| Researcher: June | -0.471 | 0.447 | -1.054 | 0.292 |
| Researcher: July | -0.432 | 0.464 | -0.932 | 0.352 |
| Researcher: August | -1.009 | 0.466 | -2.168 | **0.030** |
| Researcher: September | -1.534 | 0.548 | -2.798 | **0.005** |

**Table S10** Post-hoc comparisons of total insects observed on different crops and in different months between citizen scientist (CS) and researcher (R) collected data.

| contrast | term | estimate | SE | z.ratio | p.value |
|---|---|---|---|---|---|
| CS-R | Apple | 0.952 | 0.365 | 2.606 | **0.009** |
| CS-R | Broad bean | 1.601 | 0.313 | 5.111 | **0.000** |
| CS-R | Cherry | -0.034 | 0.528 | -0.065 | 0.948 |
| CS-R | Cucumber | 0.356 | 0.336 | 1.061 | 0.289 |
| CS-R | Currant | 1.353 | 0.390 | 3.470 | **0.001** |
| CS-R | Raspberry | 0.663 | 0.182 | 3.646 | **0.000** |
| CS-R | Runner bean | 1.602 | 0.229 | 7.001 | **0.000** |
| CS-R | Squash | 0.802 | 0.187 | 4.283 | **0.000** |
| CS-R | Strawberry | 1.582 | 0.274 | 5.779 | **0.000** |
| CS-R | Tomato | 2.697 | 0.528 | 5.109 | **0.000** |
| CS-R | April | 0.502 | 0.359 | 1.401 | 0.161 |
| CS-R | May | 0.792 | 0.194 | 4.092 | **0.001** |
| CS-R | June | 0.985 | 0.197 | 5.010 | **0.001** |
| CS-R | July | 0.955 | 0.209 | 4.574 | **0.001** |
| CS-R | August | 1.510 | 0.214 | 7.039 | **0.001** |
| CS-R | September | 1.389 | 0.374 | 3.710 | **0.001** |

**Table S11** Z-tests (with Bonferroni correction for multiple comparisons) comparing the proportion of total insect observations of each insect type between citizen scientist and researcher dataset.

| Insect Type | Z value | P value | Adjusted P value |
|---|---|---|---|
| Beetles | -1.497 | 0.134 | 0.940 |
| Hoverflies | 7.066 | 0.000 | **0.000** |
| Flies | 15.752 | 0.000 | **0.000** |
| Butterflies | 8.829 | 0.000 | **0.000** |
| Bumblebees | -8.188 | 0.000 | **0.000** |
| Honeybees | -9.436 | 0.000 | **0.000** |
| Solitary Bees | 0.200 | 0.841 | 1.000 |

**Table S12** Results of the one-way analysis of similarity (ANOSIM) comparing the similarities in crop visitation patterns between pollinator groups in data collected by the researcher (R) and data collected by citizen scientists that was matched to the researcher data by study site (CSM).

| Observer Type | Insect Type | Statistic | P Value |
|---|---|---|---|
| CSM | Butterflies | 0.561 | **0.002** |
| | Flies | 0.560 | **0.012** |
| | Wasps | 0.283 | 0.254 |
| | Hoverflies | 0.203 | 0.402 |
| R | Butterflies | 0.394 | 0.074 |
| | Flies | 0.192 | 0.430 |
| | Wasps | 0.153 | 0.514 |
| | Hoverflies | 0.120 | 0.629 |

**Table S13** Species and insect group-level metrics for the three visitation networks constructed from three data sets i) all researcher-collected data (R), ii) all citizen scientist-collected data (CS) and iii) citizen scientist-collected data matched by allotment site to the researcher-collected data (CSM).

| | Researcher Data (R) | | | Citizen Scientist Data (CS) | | | Citizen Scientists Data Matched by Site (CSM) | | |
|---|---|---|---|---|---|---|---|---|---|
| **Crop** | **d'** | **Species strength** | **N Links** | **d'** | **Species Strength** | **N Links** | **d'** | **Species Strength** | **N Links** |
| Apple/Pear | 0.086 | 0.875 | 7 | 0.054 | 0.661 | 7 | 0.052 | 0.672 | 7 |
| Broad bean | 0.169 | 0.683 | 7 | 0.061 | 0.798 | 8 | 0.153 | 0.719 | 7 |
| Cherry/Plum | 0.137 | 0.782 | 6 | 0.343 | 0.719 | 4 | 0.355 | 0.991 | 4 |
| Cucumber | 0.494 | 0.928 | 6 | 0.146 | 1.107 | 7 | 0.202 | 1.072 | 6 |
| Currants | 0.161 | 1.034 | 6 | 0.106 | 0.551 | 6 | 0.202 | 0.560 | 6 |
| Raspberry | 0.131 | 0.567 | 8 | 0.064 | 0.719 | 8 | 0.090 | 0.738 | 8 |
| Runner bean | 0.252 | 0.653 | 6 | 0.088 | 0.800 | 8 | 0.114 | 0.799 | 8 |
| Squash | 0.229 | 0.656 | 7 | 0.116 | 0.834 | 8 | 0.114 | 0.810 | 8 |
| Strawberry | 0.357 | 1.476 | 7 | 0.036 | 0.708 | 8 | 0.052 | 0.686 | 7 |
| Tomato | 0.317 | 0.345 | 2 | 0.078 | 0.872 | 8 | 0.121 | 0.954 | 7 |
| **Visitor** | | | | | | | | | |
| Beetles | 0.596 | 1.692 | 7 | 0.218 | 1.179 | 9 | 0.230 | 1.251 | 9 |
| Hoverflies | 0.214 | 0.892 | 8 | 0.115 | 1.571 | 10 | 0.145 | 1.402 | 10 |
| Flies | 0.287 | 0.984 | 8 | 0.053 | 2.661 | 10 | 0.073 | 2.766 | 10 |
| Butterflies | 0.012 | 0.047 | 5 | 0.152 | 0.561 | 8 | 0.236 | 0.495 | 7 |
| Bumblebees | 0.288 | 3.073 | 10 | 0.117 | 1.381 | 9 | 0.143 | 1.363 | 9 |
| Honeybees | 0.254 | 2.362 | 9 | 0.131 | 1.737 | 9 | 0.195 | 1.804 | 8 |
| Solitary Bees | 0.212 | 0.790 | 9 | 0.167 | 0.546 | 8 | 0.157 | 0.604 | 8 |
| Social Wasps | 0.143 | 0.164 | 6 | 0.178 | 0.365 | 9 | 0.274 | 0.315 | 7 |

**Methods S1- Survey instructions for participants**

We provided volunteers with guidance as to which crops are insect-pollinated, and participants were advised to perform their surveys on sunny, dry, and calm days where possible, ideally between the hours of 10:00-15:00. Participants were asked to record the date, time, and weather conditions at the beginning of their survey. Participants were instructed to list the name and variety, if known, of the flowering crop(s) they were surveying at that time *i.e.* those from the list of crops above that were flowering on their chosen survey date. They were asked to roughly estimate the number of plants of each crop they were surveying, using the categories provided (see Additional File S1) and to estimate the average number of open flowers per plant by examining one or two individual plants per crop. In the case of large fruit trees, participants were instructed to observe one large branch and estimate the number of open flowers on that branch.

**Methods S2- Plant-pollinator interactions analysis**

Crops were condensed into the following ten categories; apple (*Malinae* sp., apples and pears combined), broad bean (*Vicia faba*), cherry (*Prunus* sp., cherries and plums combined), cucumber (*Cucumis sativus*), currant (*Ribes* sp., redcurrant, whitecurrant, blackcurrant and gooseberries combined), raspberry (*Rubus* sp., blackberries, raspberries and loganberries combined), runner bean (*Phaseolus coccineus*), squash (*Curcubita pepo*, courgette, squash and pumpkin combined,), strawberry (*Fragaria × ananassa*) and tomato (*Solanum lycopersicum*). Crops for which there were less than fifteen observations (aubergine, gojiberry, physalis and peppers) or surveys where citizen scientists had recorded that there were no open flowers on a particular crop, were excluded from the analyses.

We used non-metric multidimensional scaling (NMDS) based on the Bray-Curtis dissimilarity index (*metaMDS* function in the "*vegan*" package (Oksanen 2019) to visualise the similarities in crop visitation between pollinator groups in data collected by the researcher and citizen scientists and tested for significant differences using a one-way analysis of similarity (ANOSIM) (*anosim* function in the "*vegan*" package). Where significant differences were observed, we used an indicator species analysis (*multipatt* function in the "*indicspecies*" package) to determine which pollinator groups were found more often in one dataset compared to the other.

We constructed a plant-visitor network from the matrix of insect pollinator counts using the "*bipartiteD3*" package (Terry 2018). We constructed three separate networks from the following data sets i) all researcher-collected data (R) ii) all citizen scientist-collected data (CS) and iii) citizen scientist-collected data matched by allotment site to the researcher-collected data (CSM). To standardise the values according to sampling effort, we followed the methods of Ballantyne et al. (2017) and calculated the visits by each insect group as a proportion of the total visits observed to a particular crop type by all insects. After visualising the networks, we calculated network indices using the "*bipartite*" package (Dormann 2022), using the same methods as in Nicholls et al. (2023).

We used $H'_2$ to estimate the level of specialisation at the level of the network. A score of zero equates to extreme generalisation and a score of one indicates perfect specialisation *i.e*. each pollinator group interacts with only one crop type. We also calculated the nestedness (weighted according to sample size) and evenness of interactions. Nestedness describes the relationship between generalist and specialist species in the network. When more specialist pollinators visit a subset of plants visited by more generalist pollinators, the network is nested (Dormann et al. 2009; Watts et al. 2016). We estimated the generality of the networks *i.e.* the mean number of partners that a plant or visitor interacts with, also weighted according to sample size (Bersier et al. 2002). At the insect/crop level, we measured the strength and specialisation (d') of interactions.
